# Synthetic Genetic Codes Designed to Hinder Evolution

**DOI:** 10.1101/695569

**Authors:** Jonathan Calles, Isaac Justice, Detravious Brinkley, Alexa Garcia, Drew Endy

## Abstract

One challenge in engineering organisms is guaranteeing system behavior over many generations. Spontaneous mutations that arise before or during use can impact heterologous genetic functions, disrupt system integration, or change organism phenotype. Here, we propose restructuring the genetic code itself such that all point mutations in protein-coding sequences are selected against. Synthetic genetic systems so-encoded should “fail safely” in response to many individual spontaneous mutations. We designed a family of such fail-safe codes and analyzed their expected effect on the evolution of engineered organisms via simulation. We predict that fail-safe codes supporting expression of 20 or 15 amino acids could slow the evolution of proteins in so-encoded organisms to 30% or 0% the rate of standard-code organisms, respectively. We also designed quadruplet-codon codes that should be capable of encoding at least 20 amino acids while ensuring all single point mutations in protein-coding sequences are selected against. We show by in vitro experiments that a reduced set of 21 tRNA is capable of expressing a protein whose coding sequence is recoded to use a fail-safe code, whereas a standard-code encoding is not expressed. Our work suggests that a rationally depleted but otherwise natural translation system should yield biological systems with intrinsically reduced evolutionary capacity, and that so-encoded hypoevolvable organisms might be less likely to invade new niches or outcompete native populations.

## INTRODUCTION

The ability to engineer organisms is increasingly important for academic, industrial, and public uses [Endy 2005; Benner and Sismour 2005; Keasling 2008; Khalil and Collins 2010; The White House 2012; Redford, Adams, and Mace 2013; Carlson 2016; Katz et al. 2018; Nye 2018]. Traditional engineering disciplines have established methods for controlling systems on the timescales of immediate input and response (e.g., data storage and retrieval, or autonomous control) [Harashima 1996, Mittal and Vetter 2016], and intermediate learning and memory (e.g., algorithms that can learn to outperform humans in games through self-play) [Silver et al. 2016, Silver et al. 2018]. However, self-reproducing systems additionally demonstrate complicated spontaneous behaviors across multiple generations [von Neumann 1966, Endy 2005]. To realize reliable operation of reproducing organisms we must also learn to engineer behavior across evolutionary timescales.

Evolution within a population relies on the diversity of genetic makeups (i.e., genotypes) from which emerges a corresponding diversity of physiological and behavioral traits (i.e., phenotypes). Genetic diversity is generated by error during DNA replication (i.e., mutation) and propagated across generations [Wright 2005, Alberts et al. 2002]. Individuals with phenotypes better suited to a given environment tend to reproduce more successfully, enriching the population with their offspring while those that are less fit are outcompeted [Alberts et al. 2002, Loewe and Hill 2010, Sniegowski and Gerrish 2010]. Thus, to engineer the evolutionary trajectories of competing populations, we must either control the processes that generate mutations or the selective pressures acting within and among populations.

One direct approach to controlling the behavior of engineered organisms over multiple generations is to reduce organism fitness outside of a prescribed niche. Scientists have long sought and realized such control of engineered organisms in safely advancing fundamental research [Berg et al. 1974, NIH 2016]. For example, biocontainment methods such as engineered auxotrophy [Ronchel and Ramos 2001, Steidler et al. 2003, Bahey-El-Din et al. 2010] or exogenously expressed “kill signals” [Gallagher et al. 2015; Callura et al. 2010; Cai et al. 2015; Agmon et al. 2017; Chan et al. 2016; Molina et al. 1998; Contreras, Molin, and Ramos 1991] have been widely used. However, such methods can be toxic to their host organisms and may result in selective pressures that inactivate the underlying mechanism [Lee et al. 2018].

More general approaches for controlling behavior over multiple generations consider altering the type and effect of mutations that arise. Such control can be realized by taking advantage of degeneracy in the mapping of DNA to proteins (i.e., the “genetic code”) to synonymously recode genes of interest [Koonin and Novozhilov 2009]. Such recoding approaches work by altering the distribution of phenotypes available to an individual without changing the identity of the translated proteins. For example, an organism can be recoded such that its initial fitness is high but nearby regions of its fitness landscape are less fit or even fatal. Such approaches have been tested by synonymously recoding Coxsackie B3 and influenza A viruses so that their genotypes were immediately adjacent to deleterious genotypes, resulting in attenuated virulence via decreased evolutionary rates [Moratorio et al. 2017]. Another approach is to recode an organism such that no single point mutation results in a significant change in fitness; organisms so-encoded might be used for a limited number of generations without fear that any single mutation will outcompete the original genotype. Such a strategy was tested by introducing infrequently used codon pairs into the poliovirus genome via synonymous recoding, resulting in both attenuated virulence and reduced likelihood of escape mutants arising during use [Coleman et al. 2008]. We note that while recoding-based approaches are generalizable to other biological systems, such approaches only affect the local fitness landscape of an organism, implying that if an engineered organism were to escape its local fitness trap it might continue to evolve unimpeded.

A more fundamental approach aims to control the entire fitness landscape of an organism by changing the underlying mapping of genotype to phenotype. Most life on Earth uses the “Standard Code” or a close variant thereof to assign 64 nucleotide triplets (i.e., “codons”) to 20 unique amino acids plus a termination signal (Fig. 1a) [Koonin and Novozhilov 2009, Hinergardner and Engelberg 1963]. The Standard Code has a highly nonrandom structure that is optimized for translation fidelity across generations (Fig. 1a-b) [Koonin and Novozhilov 2009, Koonin and Novozhilov 2017]. For example, mutations in the Standard Code are significantly more likely than in a randomly generated code to conserve the encoded amino acid (24% vs. 4%), and to minimize the physicochemical change upon mutations that do not conserve the encoded amino acid (Fig. 1d-e) [Kyte and Doolittle 1982]. Redesigning the genetic code would alter the type and effect of spontaneous mutations across all genotypes, independent of the biological system using the code. For example, recent theoretical work by Pines and colleagues proposed a “hyperevolvable” genetic code for use in directed evolution (hereafter “Colorado Code”) [Pines et al. 2017]. More specifically, Pines et al. proposed decreasing synonymous mutation rates while increasing the physicochemical changes in amino acids resulting from missense mutations (Fig. 1c-e), such that populations using the Colorado Code could traverse larger regions of phenotype space for each step in genotype space.

**Figure 1:**
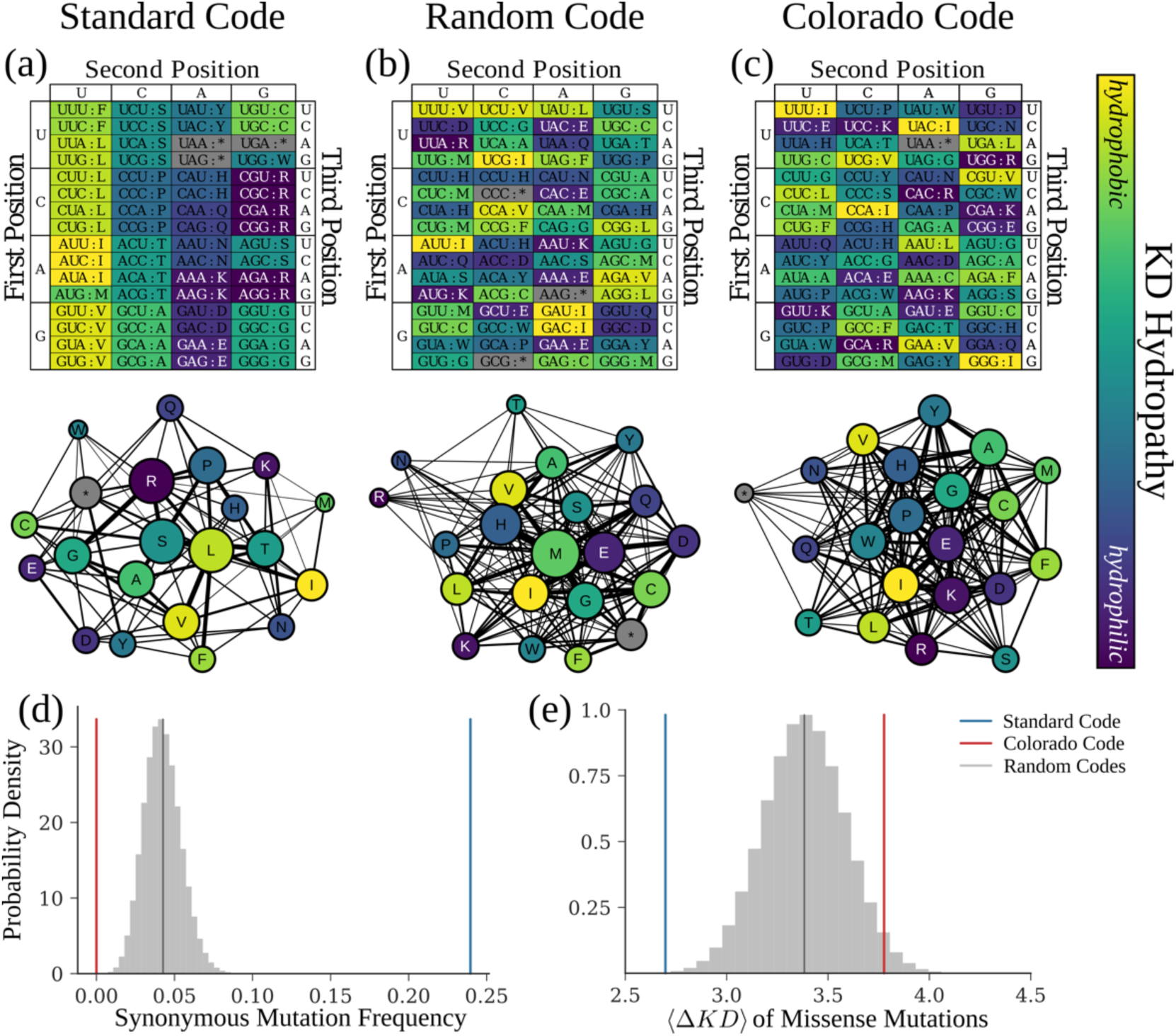
Genetic codes are expected to influence evolutionary dynamics. Table and mutation-distance network representations for (a) the Standard Genetic Code, (b) a genetic code with random structure, and (c) a theoretical hyperevolvable code from Pines et al. 2017, hereafter called the Colorado Code. For both representation styles, color signifies the rank-ordered hydropathy of the amino acids – isoleucine (I) is most hydrophobic, and arginine (R) is most hydrophilic. Mutation-distance networks represent amino acids as nodes. Node size represents the number of codons allocated to each amino acid or null. Edge weights between nodes (representing amino acids a_1_ and a_2_) represent the accessibility of a_2_ to a_1_ via point mutations. See Materials and Methods for a quantitative description of edge weighting. (d) Distribution of synonymous-mutation frequency (f_s_) for 10^6^ randomly generated codes (gray histogram) and mean of this distribution (black), as well as f_s_ for the Standard Code (blue) and Colorado Code (red). (e) Distribution of mean mutation effects given a nonsynonymous mutation (⟨ΔKD⟩) for 10^6^ randomly generated codes (gray histogram) and mean of this distribution (black), as well as ΔKD for the Standard Code (blue) and Colorado Code (red). We defined ⟨ΔKD⟩ of a genetic code as the average over all nonsynonymous mutations (from a_1_ to a_2_) of the change in Kyte-Doolittle hydropathy (ΔKD) between the two residues (ΔKD = | KD(a_2_) − KD(a_1_)|).

Here we propose a set of “fail-safe” genetic codes designed to map mutations to deleterious phenotypes, independent of the biological system in which these codes are implemented. We designed a subset of these fail-safe codes such that they might be realized using natural translation machinery, avoiding the need for molecular reengineering work. We simulated the evolutionary dynamics of populations of engineered organisms using our fail-safe genetic codes, as well as the interaction of populations using different genetic codes, in order to quantify the expected effects that different genetic codes have on evolutionary rates. We also implemented one such fail safe code using a reduced tRNA set and found that the selected reduced code is capable of synthesizing proteins *in vitro.* Our results suggest that fail-safe codes are likely to slow, or in some cases altogether arrest, the evolution of protein-coding sequences in fail-safe encoded organisms. Our results also suggest that fail-safe encoded organisms should be less able to compete with native species if introduced to new environmental contexts.

## RESULTS

### Fail-safe codes lacking translation machinery for a subset of codons are designed to penalize missense mutations

We designed a set of fail-safe genetic codes with a minimal set of translation machinery necessary to encode each expressible amino acid, eliminating degenerate sense codons (Fig. 2). Stated directly, most codons in such genetic codes are “null codons,” meaning they are not specifically recognized by any tRNAs or translation factors. Genes designed for such fail-safe codes would be encoded using the single, specific sense codon designated for each amino acid. Mutations in so-encoded open reading frames (ORFs) would most typically result in null codons. Previous work has shown that tRNA and release factor deletions that remove all machinery decoding a particular codon are either strongly deleterious or lethal, implying that translating null codons would reduce organismal fitness [Johnson et al. 2012, Bloom-Ackermann et al. 2014].

**Figure 2:**
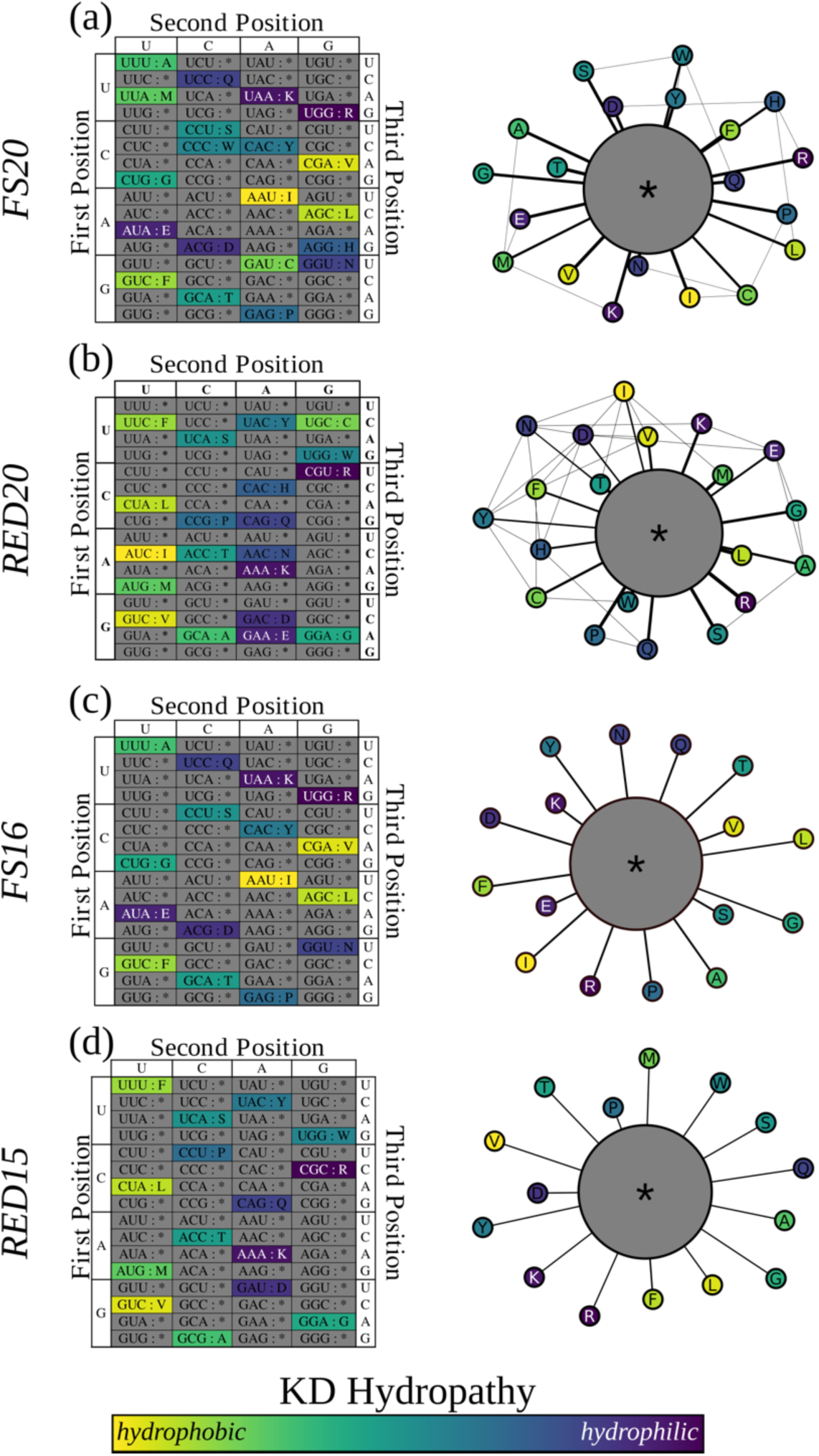
Genetic codes can be designed to map mutations from sense codons to null codons. Table and mutation-distance network representations of fail-safe codes. FS20 **(a)** requires synthetic translation machinery. RED20 **(b)** can be realized using *E. coli* translation machinery. Both FS20 and RED20 support expression with the full set of proteinogenic amino acids. FS16 **(c)** requires synthetic translation machinery. RED15 **(d)** can be realized using *E. coli* translation machinery. Both FS16 and RED15 support expression with a reduced set of amino acids such that all point mutations map to null codons. We omit specific amino acids in accordance with specific rationales (Supplementary Materials; Sup. Fig. 1).

As a first example, we designed a family of fail-safe codes in which 20 sense codons map uniquely to 20 amino acids and, to the extent possible, single point mutations map to null codons. We call these codes “Fail-Safe 20,” or FS20, because they support expression of all 20 conventional amino acids. There are P(64, 20) ≈ 5 × 10^34^ unique FS20 codes, one of which is shown as an example in Figure 2a. All FS20 codes have the same number of sense codons adjacent only to null codons, and of sense codons adjacent to each other via point mutation. However, the set of sense codons adjacent to each other via point mutation differs for each FS20 code. Engineers can therefore choose to encode their engineered organism using the FS20 code that maximizes the mutation rate to null codons given the distribution of amino acids used in the proteome of their organism. While our designs for FS20 codes anticipate eventual advances in synthetic biology sufficient to realize entirely arbitrary genetic codes, building most FW20 codes today would be nontrivial. For example, most FS20 codes would require codon reassignment involving significant reengineering of tRNAs and tRNA synthetases. While codon reassignment has been well explored for use with non-natural amino acids involving a few codons, such work has not yet been reported for all 64 codons [reviewed in Wang and Schultz 2005; reviewed in d’Aquino et al. 2018; Neumann et al. 2010; Neumann, Slusarczyk, and Chin 2010; Lajoie et al. 2013; Rovner et al. 2015; Cui et al. 2017].

To avoid reengineering all tRNAs and tRNA synthetases, we next considered synthetic genetic codes that reuse the translation machinery already implementing the Standard Code. Such genetic codes might be more readily realized by reusing naturally occurring molecules. As a first example, we designed a “reduced” fail-safe code we call RED20 (Fig. 2b, Sup. Table 1). Like FS20 codes, RED20 reduces the likelihood that mutations in protein-coding sequences result in missense mutations and increases the likelihood of mutating to a null codon. As a result, RED20 also increases the fraction of point mutations expected to result in a deleterious or lethal phenotype.

### Fail-safe codes with reduced amino acid sets or quadruplet codons only allow mutations to null codons

While FS20 and RED20 are designed to maximize the fraction of coding-sequence mutations mapping to null codons and minimize the fraction of missense mutations, it is impossible to encode 20 amino acids in a 64-codon genetic code such that each sense codon is only immediately adjacent to null codons. Eliminating all missense mutations in a genetic code and ensuring that all mutations from sense codons map to null codons required we consider either encoding fewer amino acids or adopting a larger codon table.

Thus, we designed a family of fail-safe codes based on the FS20 codes that encode reduced sets of 16 amino acids (hereafter FS16, Fig. 2c). FS16 codes are designed such that no single sense codon can mutate to any other sense codon via a single point mutation. Similarly, we designed a fail-safe code based on RED20 that encodes 15 amino acids, mapping all mutations to null codons, that can be built via naturally occurring translation machinery alone (hereafter RED15, Fig. 2d). Because FS16 and RED15 map all mutations to null codons we call them “ideal” fail-safe codes. We selected and recommend specific FS16 and RED15 codes on the basis of various design principles (e.g., if one of many similar amino acids is encoded then other similar amino acids become less important) (Supplementary Materials; Sup. Fig. 1).

We also considered genetic codes with expanded codon sets. Quadruplet decoding occurs in nature [Gesteland, Weiss, and Atkins 1992] and has been demonstrated experimentally [Magliery, Anderson, and Schultz 2001; Neumann et al. 2010; Niu et al. 2013; Wang et al. 2012]. While the use of quadruplet codons is currently limited to a few positions per gene [Neumann et al. 2010; Niu et al. 2013; reviewed in Wang et al. 2012], we considered quadruplet codon designs in anticipation of ongoing advances in synthetic biology. Specifically, we designed a family of quadruplet-codon fail-safe codes (hereafter FSQUAD) with 256 available codons (Sup. Fig. 2). FSQUAD codes would be able to encode more than 20 amino acids such that all mutations from sense codons map to null codons, allowing for programmable incorporation of non-natural amino acids in a fail-safe encoded system. Like FS20- or FS16-encoded organisms, an FSQUAD-encoded organism should also be resistant to horizontal gene transfer.

### Simulations quantify relative evolutionary rates of different genetic codes

To predict how fail-safe genetic codes affect evolution we simulated large, asexual populations of organisms encoded via fail-safe genetic codes. We developed a hybrid model where small population-size lineages are treated stochastically using a birth-death process to capture genetic drift, and large population-size lineages are treated deterministically with exponential growth. Mutations are generated stochastically, the number of which is dependent on the population size and genetic code used [Desai and Fisher 2007; Desai, Fisher, and Murray 2007]. During the course of a simulation an initially monoclonal population generates diversity via mutation. Newer, more fit strains arise and slowly outcompete less fit strains, increasing the mean fitness of the population (Fig. 3a). We compare the evolutionary rate of genetic codes by comparing the different rates of increasing fitness across populations using these codes (Fig. 3b). For example, in the systems being studied we predict the Standard Code allows fitness to increase at a rate of 8.71 × 10^−4^ 1/gen^2^(s.d 1.31 × 10^−4^ 1/gen^2^). In comparison, the Colorado Code is expected to evolve only slightly faster (9.79 × 10^−4^ 1/gen^2^, s.d. 1.36 × 10^−4^ 1/gen^2^, or 12.4% faster).

**Figure 3:**
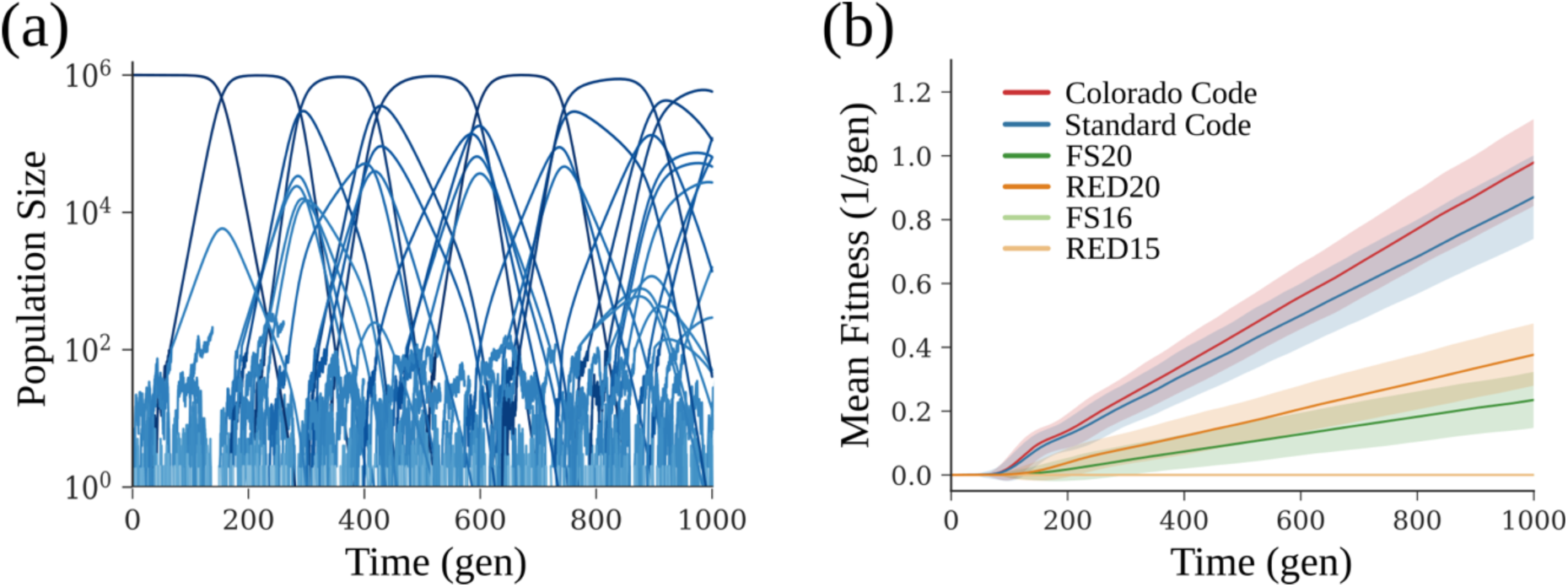
Simulations suggest fail-safe codes can attenuate evolution more effectively than hyperevolvable codes can accelerate evolution. **(a)** A simulation of mutation-selection balance in large, asexual populations. Each line represents the population size of an isogenic lineage in the simulation vs. time. New lineages arise as mutants are generated. A detailed description of the model is given in the Materials and Methods section. **(b)** Mean fitness traces for replicates of populations (n = 1000) using the Standard Code (blue), Colorado Code (red), FS20 (dark green), FS16 (light green), RED20 (dark orange), and RED15 (light orange). Bold lines indicate the mean fitness of a batch culture, averaged across replicates. Shaded regions represent the standard deviation across replicates.

The fail-safe codes studied here are expected to have a much stronger effect on evolutionary dynamics (Table 1). For example, FS20 reduced predicted evolutionary rates to 27% that of the Standard Code (2.35 × 10^−4^ 1/gen^2^, s.d. 0.877 × 10^−4^ 1/gen^2^). RED20 behaves qualitatively similarly to FS20, despite its imposed design constraints, yielding an expected evolutionary rate only 43% that of the Standard Code (3.77 × 10^−4^/gen^2^, s.d. 0.977 × 10^−4^ 1/gen^2^). The ideal fail-safe codes FF16 and RED15 were predicted to arrest ORF evolution due to single point mutations altogether, and thereby maintaining their initial population fitness over the duration of the simulation.

**Table 1:**
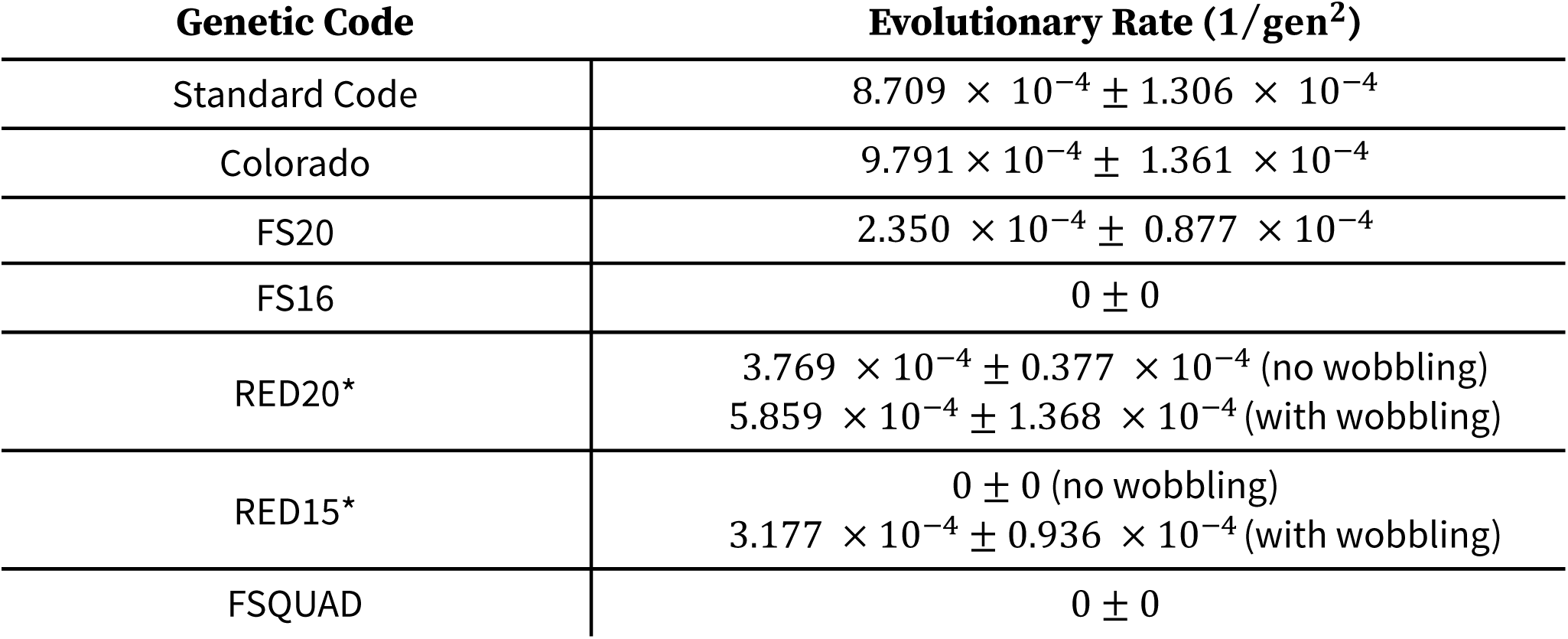
Evolutionary rates are expected to vary across natural and fail-safe genetic codes. Summary of evolutionary dynamics simulations. Predicted evolutionary rate is reported as the change in fitness (in units of 1 */ gen*) per unit time (in units of *gen*). Mean rate of fitness increase is reported along with standard deviation. Codes marked with an asterisk were simulated both with and without considering tRNA promiscuity.

| Genetic Code | Evolutionary Rate (1/gen<sup>2</sup>) |
| --- | --- |
| Standard Code | $8.709 \times 10^{-4} \pm 1.306 \times 10^{-4}$ |
| Colorado | $9.791 \times 10^{-4} \pm 1.361 \times 10^{-4}$ |
| FS20 | $2.350 \times 10^{-4} \pm 0.877 \times 10^{-4}$ |
| FS16 | $0 \pm 0$ |
| RED20* | $3.769 \times 10^{-4} \pm 0.377 \times 10^{-4}$ (no wobbling)<br>$5.859 \times 10^{-4} \pm 1.368 \times 10^{-4}$ (with wobbling) |
| RED15* | $0 \pm 0$ (no wobbling)<br>$3.177 \times 10^{-4} \pm 0.936 \times 10^{-4}$ (with wobbling) |
| FSQUAD | $0 \pm 0$ |

### Biocontainment may arise intrinsically in organisms using fail-safe genetic codes

We hypothesized that fail-safe encoded organisms will adapt to new environments more slowly than naturally encoded organisms and thus might be less able to challenge established, native populations. If true then fail-safe encoding could be used as an intrinsic biocontainment layer, one that does not rely on a heterologous genetic function but rather is instantiated via the encoding of the entire organism. To quantitatively assess this possibility, we simulated competing populations of organisms encoded by both Standard and fail-safe codes, exploring when and to what extent invading populations might displace established populations. In our simulations the invasive populations either swept or were swept by the native populations (Fig. 4a). More specifically, we defined a containment probability, P_contain_(f_0_, t, 𝕋), as the likelihood that the invasive population will have been outcompeted by time t, given an initial invasive population fraction f_0_ and genetic code 𝕋. After sufficient time P_contain_ reaches a steady state, varying only in initial population fraction (Fig. 4b, Sup. Fig. 5). We generated approximate steady state containment curves (Fig. 4c). We predict FS20 will maintain a containment probability P_contain_ < 99% up to an initial invasive population fraction f_0_ ≤ 36%. RED20 was able to maintain P_contain_ < 99% up to f_0_ ≤ 14%. We predict organisms encoded in FS16 and RED15 would be outcompeted across all initial conditions simulated. Our results suggest that population-level biocontainment is expected to be an intrinsic property of organisms encoded via fail-safe codes.

**Figure 4:**
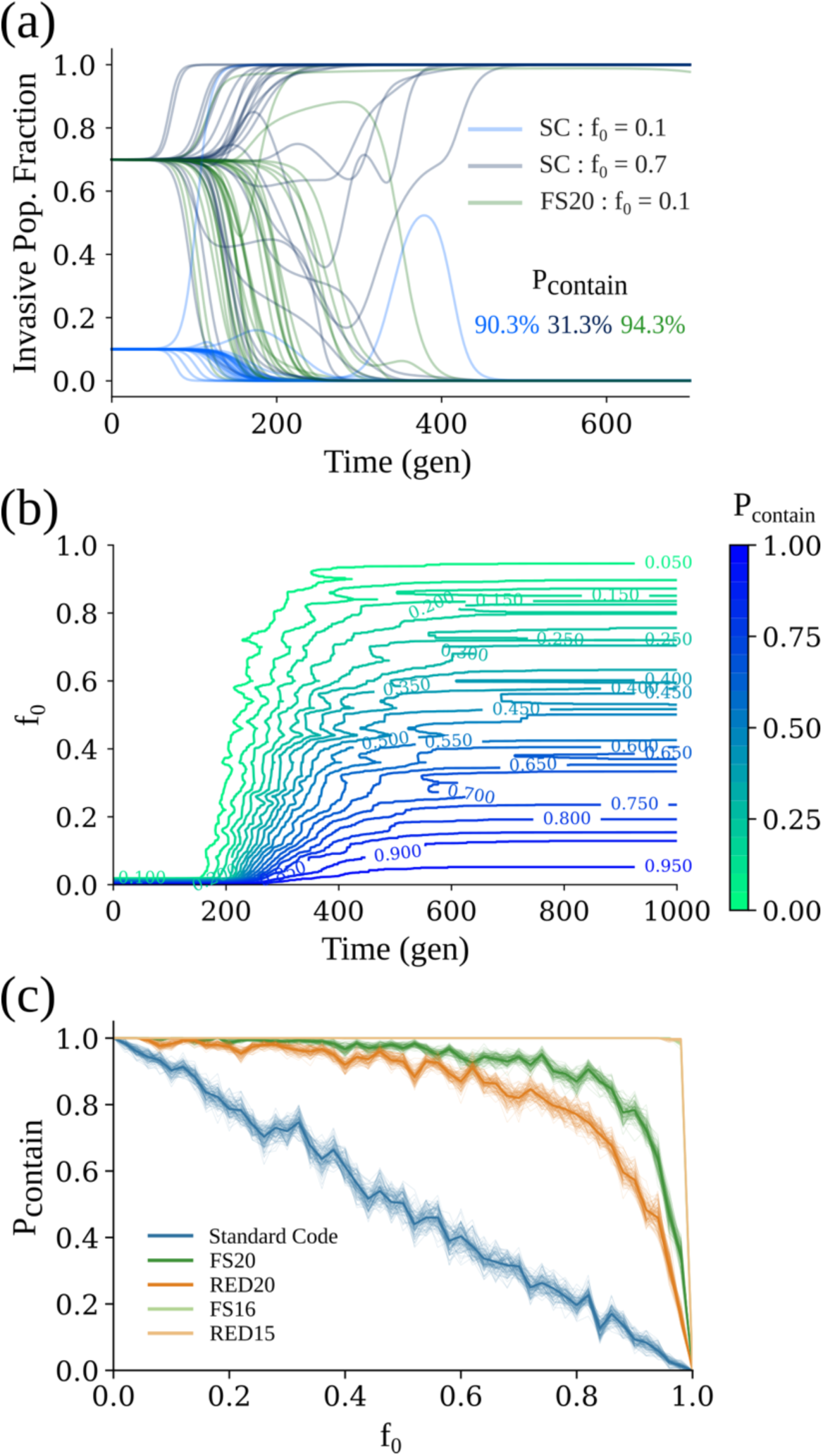
Fail-safe codes may also prevent organisms from escaping into the environment. **(a)** Replicates (n = 300) of simulated competition between a native population encoded in the Standard Code and a monoclonal invasive population either encoded in the Standard Code with an initial population fraction *f*_0_ = 10% (light blue) or 70% (dark blue), or encoded in FS20 with *f*_0_ = 70% (green). We approximate containment probability *P*_*contain*_ as the fraction of simulations in which the invasive population is eliminated. **(b)** Contour graphs of containment probability vs. time (x axis) and *f*_0_ (y axis) for invasive strains using the Standard Code (n = 300 replicates). Color represents *P*_*contain*_ magnitude, varying from 0 (green) to 1 (blue). *P*_*contain*_ reaches a steady state value at the limit of large *t*. **(c)** *P*_*contain*_ at steady state vs. for invasive strains using fail-safe codes (n = 300 replicates). Bootstrapped-resampled traces of the data are represented as lighter shaded lines. Colors are the same as in Fig. 3b.

**Figure 5:**
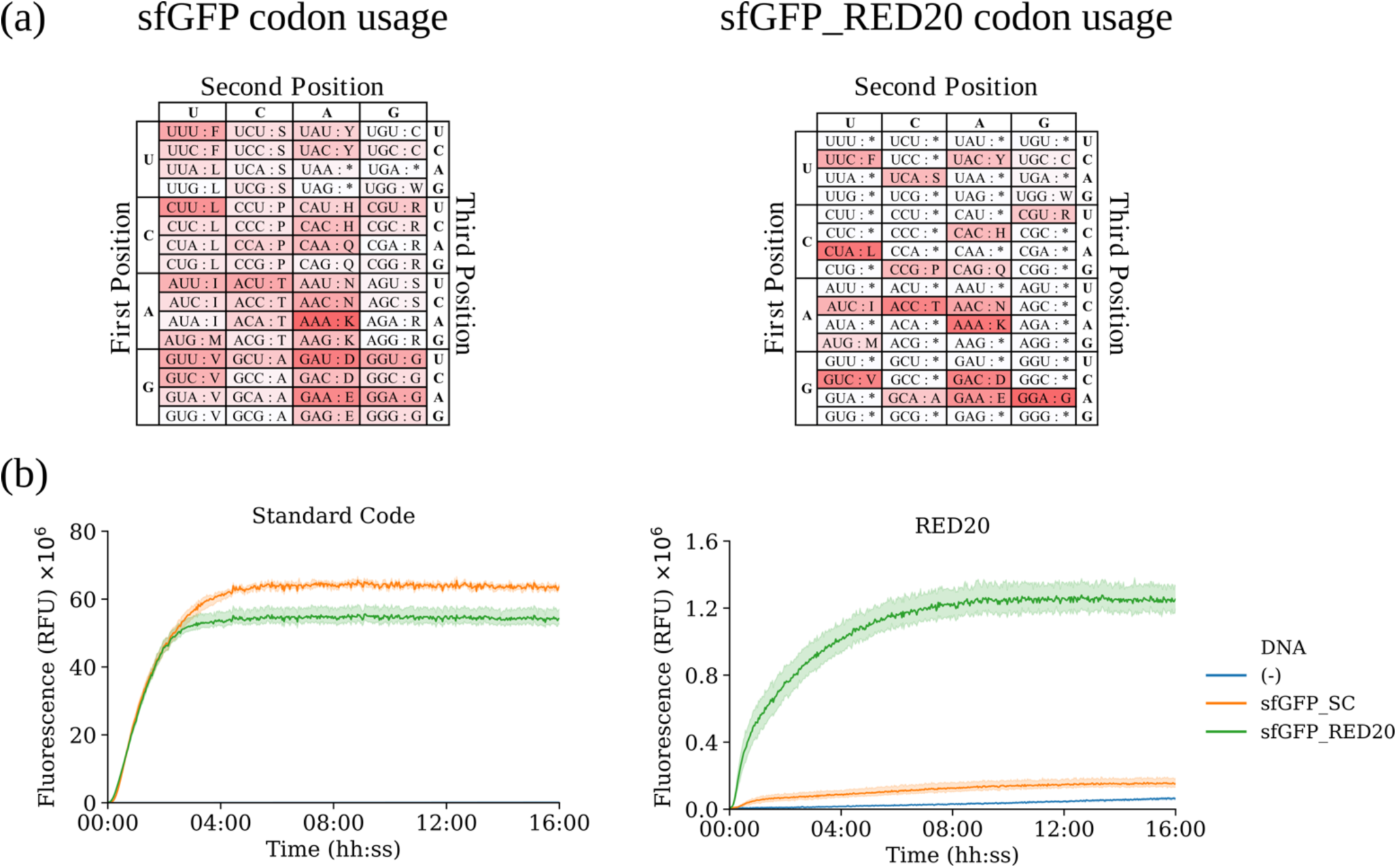
A reduced set of tRNA encoding RED20 can express a functional fluorescent protein. **(a)** Frequency of codon usage in coding sequences for super folder variants of GFP (sfGFP) encoded in either the Standard Code (sfGFP_SC, left) or RED20 (sfGFP_RED20, right). Unused codons are represented in white, while frequently used codons are represented in red. **(b)** Fluorescence versus time for sfGFP variants expressed *in vitro* from a tRNA set encoding either the Standard Code (left) or RED20 (right). Blue lines represent negative control traces (without reporter DNA, n=3). Orange and green regions represent fluorescence traces from sfGFP encoded in the Standard Code (n=3) and RED20 (n=4), respectively, with mean values shown as bold traces.

### A reduced set of tRNAs instantiating RED20 enables protein expression

We sought to prototype a translation system using one of our fail-safe codes to learn if our any of our designs might have a chance of working. PURE is a chemically defined *in vitro* translation system composed of individually purified components [Shimizu et al. 2001]. We obtained PURE lacking all native tRNAs (PURE ΔtRNAs). We also procured a reduced set of tRNA instantiating RED20 via direct RNA synthesis, which we added to PURE ΔtRNAs to make an *in vitro* RED20 expression system. We recoded green fluorescence protein to use only the RED20 codons (Fig. 5a). We found that our system using only RED20 tRNA is able to successfully express RED20-encoded, but not standard-encoded, fluorescent protein. Specifically, we observed that our prototype RED20 system expressed RED20-encoded sfGFP at a level 8-fold higher than standard-encoded sfGFP (Fig. 5b).

## DISCUSSION

We designed fail-safe genetic codes that lack translation machinery recognizing the majority of codons such that individual point mutations in protein coding sequences should be deleterious to the host organism. We then simulated the evolution of populations using these codes to quantitatively predict the expected effects of fail-safe genetic codes on evolutionary dynamics. Our designed codes were able to reduce evolutionary rate to ∼30% of the Standard Code while encoding the full set of 20 conventional amino acids and to select against all individual point mutations in coding sequences if encoding only 15 or 16 amino acids. The most practical-to-implement codes, RED15 and RED20, are predicted to behave qualitatively similarly to FS16 and FS20 respectively without requiring any tRNA and tRNA synthetase engineering. Further, we built a functional RED20 prototype *in vitro* and demonstrated its capacity for protein expression.

### Fail-safe codes may serve as a base layer for biocontainment strategies

Previous work has focused on containing organisms to prescribed physical niches [Steidler et al. 2003; Ronchel and Ramos 2001; Bahey-El-Din et al. 2010; Gallagher et al. 2015; Callura et al. 2010; Cai et al. 2015; Agmon et al. 2017; Chan et al. 2016; Molina et al. 1998; Contreras, Molin, and Ramos 1991]. However, full control of reproducing populations will also require containing organisms to prescribed genotypes. To ensure the reliability and long-term stability of synthetic genetic programs we need “genetic containment” methods. Our work suggests that fail-safe codes can offer both physical and genetic containment. Specifically, we predict that fail-safe encoded organisms will not only explore genotype space slower than organisms encoded using the Standard Code but will also be less likely to outcompete native populations in new environmental contexts. Organisms encoded with fail-safe codes such as FS20 or FS16 would additionally be genetically isolated from natural organisms [Lajoie et al. 2013, Ravikumar and Liu 2015]. We believe that fail-safe codes can be used as a base containment layer upon which additional safeguards can be added modularly [Gallager et al. 2015].

### Wobble decoding presents a general challenge for code engineering

One challenge in code engineering is the tendency for tRNAs to recognize more than one codon due to wobble decoding [Crick 1966; Tuite 2001; Agris, Vendeix, and Graham 2007; Watanabe and Osawa 1995] For example, designs for a hyperevolvable code generally maximize the diversity of encoded amino acids adjacent to any given sense codon, which can result in an ambiguous code where many codons are recognized by two differentially aminoacylated tRNAs (Sup. Fig. 3). The effect of wobble decoding on fail-safe codes however is comparatively less drastic. We simulated the behavior of organisms using RED20 and RED15 assuming 100% efficient wobble decoding (Sup. Fig. 4). Under these assumptions, we predict an evolutionary rate for RED20 and RED15 of 65% and 37% of the Standard Code rate, respectively. We further predict under these assumptions that organisms using RED20 and RED15 maintain a containment probability greater than 95% up to an invading population fraction (f_0_) of 23% and 54%, respectively. Therefore, while engineering one-to-one decoding would improve the efficacy of fail-safe codes, we predict that RED15 and RED20 are robust to wobble decoding even if instantiated via native or near-native tRNA.

Predicting how wobble decoding might affect a quadruplet code is difficult. We may naively assume that the additional base pair in the codon-anticodon complex would allow FSQUAD to encode four times as many amino acids unambiguously. If so, an ideal quadruplet fail-safe code may be able to encode up to 32 sense positions adjacent only to null codons without requiring tRNAs capable of one-to-one decoding. However, engineering a full set of quadruplet decoding tRNAs, the cognate aminoacyl transferases and translation factors, and maintaining perfect codon-anticodon specificity would be challenging.

### Reduced amino acid sets may still encode interesting biological functions

One way to increase the probability of mutating to a null codon in a fail-safe code is to decrease the number of encoded amino acids, thereby decreasing the number of required sense codons. But what biological functions can be encoded with less than 20 amino acids? Could a whole organism ever be encoded with a reduced amino acid set? Of the 20 proteinogenic amino acids, ten are predicted to have resulted from biosynthesis in early organisms [Miller 1953; Danger, Plasson, and Pascal 2012; Fujishima et al. 2018]. This implies relevant biological functions may have been encoded with as few as ten amino acids. As a first step towards a reduced amino acid set organism, we recently removed cysteine from all enzymes in the cysteine biosynthesis pathway [Fujishima et al. 2018]. However, significant additional work would be required to remove four or five amino acids starting from any known natural organism, as would be needed to realize a FS16 or RED15 code.

Additionally, we searched the UniProt database [The UniProt Consortium 2017] to see if any existing natural proteins use less than 20 amino acids. As one example, we found the antimicrobial peptide acanthoscurrin-2 is encoded only via amino acids in our RED15 code [Lorenzini et al. 2003]. We also analyzed residue conservation in the green fluorescent protein (GFP) using ConSurf [Ashkenazy et al. 2016] to assess which residues are most essential to protein function and what diversity of amino acids are found at these positions, finding that it should be possible to realize a functional GFP using only the RED15 translation system (Sup. Table 2, Sup. Fig. 6).

### Gene duplication and tRNA evolution are expected failure modes of fail-safe codes

We expect that increasing the rate of mutations to null codons will add a selective pressure for noncognate translation machinery to recognize these null codons. For example, ribosomal ambiguity mutations (ram) impair the proofreading ability of the ribosome [Gorini, Jacoby, and Breckenrirdge 1966; Rosset and Gorini 1969], increasing the likelihood that a noncognate tRNA can recognize a null codon. While several ram mutations have been discovered in ribosomal proteins [Piepersberg, Böck, and Wittmann 1975; Cabezón et al. 1976; Kirsebom and Isaksson 1985; Agarwal et al. 2011; Agarwal et al. 2015], we expect that ram mutations in rRNA [McClory et al. 2010; McClory et al. 2014; Santer et al. 1995; O’Connor et al. 1995; O’Connor et al. 1997; Gregory, Lieberman, and Dahlberg 1994; Murgola et al. 1995] are more likely to accumulate in a fail-safe encoded organism given such genetic codes only affect mutations in protein-coding sequences. We also note that fail-safe codes do not prevent gene duplication. Chromosomal and whole-genome duplication events can result in novel genetic functions [Kasahara et al. 1996, Wolfe and Shields 1997; De Bodt, Maere, and Van de Peer 2005], frequently as a response to stress [Yona et al. 2012]. Duplication of tRNA genes specifically and subsequent mutation of the anticodon loop has been suggested as a mechanism for genetic code reprogramming in nature [Schultz and Yarus 1994, Osawa and Jukes 1995]. Such a mechanism could generate tRNAs that recognize null codons, subverting an evolutionary containment strategy based on a fail-safe code. Duplication and subsequent mutation as an evolutionary mechanism has been experimentally validated in other contexts (e.g., *E. coli* lactose metabolism [Kugelberg et al. 2006]). Such failure modes, and likely others, would need to be addressed in order to realize fully nonevolving organisms.

### Removing sense codons from a genome presents a technical challenge

Building a fail-safe encoded organism will require the ability to encode an entire genome such that each amino acid is represented by only one codon. However, codon usage has been shown to regulate gene expression, translation speed, and co-translational folding of proteins [Hershberg and Petrov 2008, Buhr et al. 2016, Escudero et al. 2017], as well as affect the overall fitness of the organism [Coleman et al. 2008]. As a result, some synonymous codon substitutions appear disallowed *in vivo* [Wang et al. 2016]. It is an open question how many sense codons are required to instantiate a living organism. Recently, Fredens and colleagues created a synthetic variant of the *E. coli* genome using only 61 codons, 59 of which encode amino acids via synonymous recoding of 18,214 codons plus deletion of otherwise-essential tRNA [Fredens *et al.* 2019]. Additionally, Ostrov and colleagues are working to remove seven sense codons from *E. coli* to create a 57-codon organism, and have reported successfully recoding 60% of *E. coli* genes [Ostrov et al. 2016]. While both examples demonstrate the state-of-art in genome-scale codon reduction, they fall short of the scope of reduction necessary to realize a 15 or 20 sense-codon organism. However, recently developed tools that accelerate total genome synthesis have enabled researchers to more rapidly screen recoding strategies, accelerating the pace of progress in the field of codon reduction [Wang et al. 2016]. As engineering whole genomes becomes more feasible so too should designing and building genomes with fewer and fewer sense codons.

We believe that fail-safe codes will play an important foundational role in controlling the evolution of biological systems, especially in the context of whole genome engineering. We noted several challenges that need to be addressed before the first fail-safe organism can be realized. Importantly, a subset of our proposed codes do not require reassigning sense codons, relying instead only on the removal of some isoacceptor tRNAs from the natural translation system, greatly simplifying initial experiments. Given the importance of exploring and realizing non-evolving biological systems, as well as the practicality of validation experiments, we hope that additional academic work on fail-safe codes will be quickly complemented by public and private efforts to realize fail safe organisms providing a best available technology for realizing responsible engineered organisms suitable for deployment in field, plant, animal, or patient.

## MATERIALS AND METHODS

### Software

All code used herein is free online via https://github.com/EndyLab/codon-tables/tree/manuscript

### Constructing mutation-distance networks

We made abstract visualizations of the genetic codes considered in this work to understand the single and multiple point mutations available to any given code. We converted genetic codes into force-directed graphs. Nodes represent encoded amino acids and edges represent mutations between sense codons. Two nodes are connected by an edge if there exists at least one pair of codons (c_1_ and c_2_) encoding amino acids (a_1_ and a_2_) such that c_1_ can be converted to c_2_ by a single point mutation. The edge weight between any two amino acids a_1_ and a_2_ takes into account all possible acyclic paths between the set of codons encoding a_1_ and a_2_, respectively, including indirect paths that involve initial synonymous mutations. Individual paths from c_1_ to c_2_ are weighted by an inverse power law representing the number of point mutations necessary to convert c_1_ to c_2_. Paths are then summed to give the total edge weight.

Formally, let C = {UUU, …, GGG} be the set of all triplet codons, A = {F, L, …, G} be the set of all amino acids, T: c → a | c ∈ C, a ∈ A be a genetic code, and W(a_1_, a_2_) be the edge weight between amino acids a_1_ and a_2_:

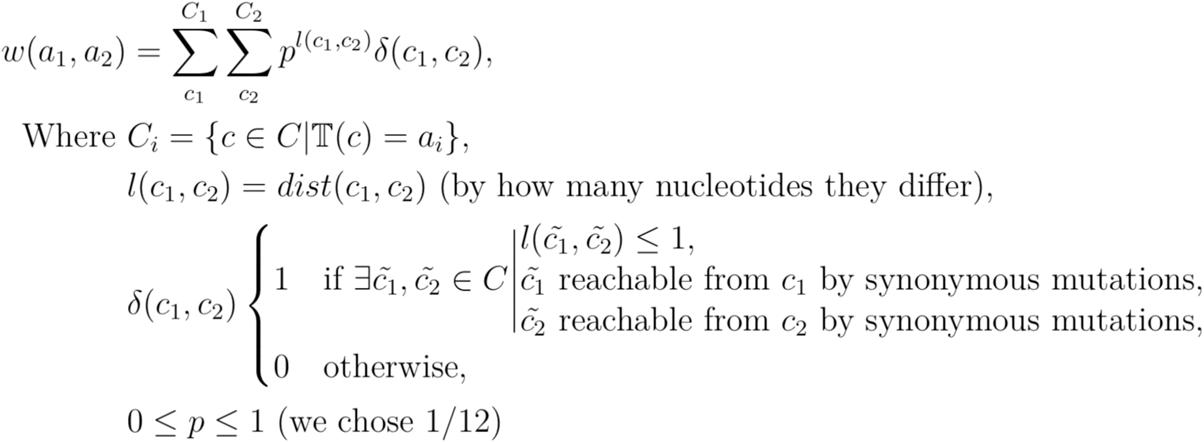

### Modeling wobble decoding and tRNA promiscuity

Here, we chose to represent sense codon decoding using the most specific tRNA species for a given codo (i.e., the tRNA recognizing the fewest additional codons). We used the following heuristic rules: NNY codons (with U or C in the wobble position) can be decoded by anticodons GNN and QNN (where Q is queuosine), however generally tRNAs cannot discriminate NNU from NNC. Similarly, NNR (with A or G in the wobble position) are decoded by tRNAs with modified uridine in the 34th position, e.g., cmnm5U, mcm5U, Um, and xm5s2U. While Ile-tRNACAU can distinguish AUA from AUG using k2C in the 34th position, this ability to decode NNA and not NNG does not generalize to all NNA decoding species. NNG is fully distinguishable from NNH (all but NNG) codons simply with an unmodified C in the 34th position [Yokoyama and Nishimura 1995, Watanabe and Osawa 1995]. Please see Sup. Table 1 for a description of the RNA base modification shorthand used herein.

### Simulating evolutionary dynamics

All simulations were carried out in Python 3.6.4 on Docker instances (running Debian 8) hosted by Amazon Web Services (AWS). Parallelization was managed by AWS Batch. Each simulated strain was partitioned into one of two groups based on population size, which are modeled independently over a small epoch dt (chosen at 0.1 gen). We modeled small population-size group using a stochastic birth-death model. The per-individual doubling probability in an epoch is given by p_b_ = [1 + (f_i_ − ⟨f⟩)]dt where f_i_ is strain’s fitness and ⟨f⟩ is the mean fitness of the batch culture. The corresponding death probability is fixed at p_d_ = (1)dt. We modeled the large population-size group analytically with strain size N_i_ given by 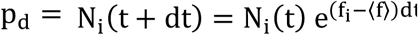. At the end of each epoch, we recalculated the mean fitness of the simulated batch culture and reallocated strains between the low and high population-size groups. The threshold population size at which a strain is reallocated (ϵ_i_) is strain specific and given by 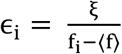, where ξ is a constant factor (we chose ξ = 3).

We modeled mutation and the generation of new strains in two steps. We first determine the number of mutants each strain will generate in a given epoch by drawing from a Poisson distribution with expectation value for each strain µ_i_ = N_i_U_b_ϕ_i_dt, where N_i_ is the strain’s population size, U_b_ is the per genome per generation beneficial mutation rate (set at 10^−5.5^), and dt is the epoch duration. ϕ_i_ is calculated as the fraction of missense mutations in a genetic code that do not result in truncation, normalized by that same fraction for the Standard Code.

Each mutation is then assigned a fitness effect (df_i_), drawn from a Distribution of Fitness Effects (DFE). We modeled the DFE with a generalized half-normal distribution 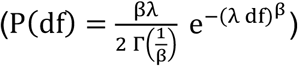. We then introduce a new strain for each mutation, with population size N_i_ = 1 and fitness f_i_ = f_j_ + df_i_, where f_j_ is the fitness of the parent strain from which the new strain mutated.

We made two approximations to reduce computational load. Our first approximation relies on the theoretical result that mutants generated from strains with low population-sizes have a vanishingly low probability of establishing in the population [Desai and Fisher 2007]. Thus, to reduce computational load, we did not generate mutants originating from the small population-size group. Our second approximation prematurely removes low fitness strains from the population once two conditions are met: (1) strain fitness has dropped below the mean, and (2) after strain population size has reduced such that the strain would be moved to the small population-size group and modelled stochastically. While this artificially inflates the mean fitness of the simulated batch culture, the effect is small given the small population-size’s total contribution to the weighted average of fitness of the batch culture (on the order of 0.03%). We observed that these approximations greatly improved simulation speed without qualitatively affecting the results.

### Preparing expression plasmids

We received pSB1C3-T7-sfGFP from Eric Wei as a gift, which we used as the Standard Code-encoded expression vector (sfGFP_SC), as well as the backbone of our RED20-encoded expression vector (sfGFP_RED20). To produce sfGFP_RED20, we first computationally recoded the sfGFP coding sequence to only include codons used by RED20. The recoded gene was then synthesized *ab initio* by Twist Biosciences and assembled into pSB1C3-T7 using the NEB HiFi Assembly kit (NEB# E5520S).

Chemically competent *E. coli* Top10 cells were incubated with 2.5 µL of assembly product on ice for 30 minutes. These cells were then heat shocked at 42C for 30s, returned to ice for two minutes, then grown out in 950 µL SOC media at 37C for one hour. The resulting transformants were plated on LB agar with chloramphenicol (25 ng/µL) and grown over night at 37C. Colonies were then grown up in 50 mL TB broth with chloramphenicol (25 ng/µL) for 16 hours. Each overnight culture was split into five batches of 10 mL each, and plasmid was prepped from each batch separately using QIAprep Spin Miniprep kits (QIAGEN, Cat No./ID: 27104) and then pooled. Final DNA product was assessed for quantity and purity using a NanoDrop 2000 (Thermo Scientific). Annotated sequence maps for sfGFP_SC (https://benchling.com/s/seq-gqXNUQJ41NbxOmdFD3LN) and for sfGFP_RED20 (https://benchling.com/s/seq-w63RBxrXRxi6uIruvKEM) can be found online.

### Expressing protein and measuring fluorescence in vitro

Twenty-one tRNA species were chemically synthesized *ab initio* by Agilent Technologies and resuspended in nuclease free TE buffer at pH 8.0. These tRNAs were then combined in equimolar ratio, at 250 mM each, to create a RED20 tRNA 25x master mix (10 mM final concentration per tRNA). An *in vitro* RED20 prototype was prepared by supplementing PURExpress *in vitro* expression system lacking tRNAs (PURE ΔtRNA, NEB# E6840S). PURE ΔtRNA supplemented with supplied control tRNAs was used as a Standard Code. We added 1 µL of murine RNase inhibitor to all *in vitro* reaction (NEB# M0314S). Each reaction also received 60 pmol of either the RED20-encoded or the standard-encoded expression vector. Otherwise, reactions were assembled as specified by NEB to a final volume of 10 µL. Reactions were carried out in a SpectraMax i3 plate reader (Molecular Devices) using clear bottom, 384-well microtiter plates (Corning) at 37C for 16 hours. Protein expression was measured using the same plate reader. Samples were excited at 485 nm (9 nm bandwidth), and emission was measured at 520 nm (15 nm bandwidth). At two minute read intervals samples were read from the bottom of the plate following three seconds of shaking prior to each measurement.

## ACKNOWLEDGEMENTS

The authors would like to thank Akshay Maheshwari and Sam Bray for helpful discussions regarding principles and applications of genetic code engineering and controlling evolution, Conary Meyer, Eric Wei, Rolando Perez and Keoni Gandall for helpful conversations regarding physical implementation of these proposed genetic codes, Anton Jackson-Smith for substantial contributions to the codebase of this project, both in advising us on structuring the simulation code as well as for writing components of our parallelization and bioinformatic analysis scripts, and Austin Che along with the OpenWetWare community for long-ago conversations regarding fail-safe genetic codes.

## REFERENCES

1. Agarwal, D., Gregory, S. T. & O’Connor, M. Error-prone and error-restrictive mutations affecting ribosomal protein S12. J. Mol. Biol. 410, 1–9 (2011).

2. Agarwal, D., Kamath, D., Gregory, S. T. & O’Connor, M. Modulation of decoding fidelity by ribosomal proteins S4 and S5. J. Bacteriol. 197, 1017–25 (2015).

3. Agmon, N. et al. Low escape-rate genome safeguards with minimal molecular perturbation of Saccharomyces cerevisiae. Proc. Natl. Acad. Sci. U. S. A. 114, E1470–E1479 (2017).

4. Agris, P. F., Vendeix, F. A. P. & Graham, W. D. tRNA’s Wobble Decoding of the Genome: 40 Years of Modification. J. Mol. Biol. 366, 1–13 (2007).

5. Alberts, B. et al. Molecular Biology of the Cell. (Garland Science, 2002).

6. Ashkenazy, H. et al. ConSurf 2016: an improved methodology to estimate and visualize evolutionary conservation in macromolecules. Nucleic Acids Res. 44, W344–W350 (2016).

7. Bahey-El-Din, M., Casey, P. G., Griffin, B. T. & Gahan, C. G. Efficacy of a Lactococcus lactis ΔpyrG vaccine delivery platform expressing chromosomally integrated hly from Listeria monocytogenes. Bioeng. Bugs 1, 66–74 (2010).

8. Bateman, A. et al. UniProt: The universal protein knowledgebase. Nucleic Acids Res. 45, D158–D169 (2017).

9. Benner, S. A. & Sismour, A. M. Synthetic biology. Nat. Rev. Genet. 6, 533–543 (2005).

10. Berg, P. et al. Potential biohazards of recombinant DNA molecules. Science 185, 303 (1974).

11. Bloom-Ackermann, Z. et al. A Comprehensive tRNA Deletion Library Unravels the Genetic Architecture of the tRNA Pool. PLoS Genet. 10, e1004084 (2014).

12. Buhr, F. et al. Synonymous Codons Direct Cotranslational Folding toward Different Protein Conformations. Mol. Cell 61, 341–351 (2016).

13. Cabezón, T., Herzog, A., De Wilde, M., Villarroel, R. & Bollen, A. Cooperative control of translational fidelity by ribosomal proteins in Escherichia coli. Mol. Gen. Genet. MGG 144, 59–62 (1976).

14. Cai, Y. et al. Intrinsic biocontainment: Multiplex genome safeguards combine transcriptional and recombinational control of essential yeast genes. Proc. Natl. Acad. Sci. 112, 1803–1808 (2015).

15. Callura, J. M., Dwyer, D. J., Isaacs, F. J., Cantor, C. R. & Collins, J. J. Tracking, tuning, and terminating microbial physiology using synthetic riboregulators. Proc. Natl. Acad. Sci. U. S. A. 107, 15898–903 (2010).

16. Carlson, R. Estimating the biotech sector’s contribution to the US economy. Nat. Biotechnol. 34, 247–255 (2016).

17. Chan, C. T. Y., Lee, J. W., Cameron, D. E., Bashor, C. J. & Collins, J. J. ‘Deadman’ and ‘Passcode’ microbial kill switches for bacterial containment. Nat. Chem. Biol. 12, 82–86 (2016).

18. Coleman, J. R. et al. Virus Attenuation by Genome-Scale Changes in Codon Pair Bias. Science (80-.). 320, 1784–1787 (2008).

19. Contreras, A., Molin, S. & Ramos, J. L. Conditional-suicide containment system for bacteria which mineralize aromatics. Appl. Environ. Microbiol. 57, 1504–8 (1991).

20. Crick, F. H. Codon--anticodon pairing: the wobble hypothesis. J. Mol. Biol. 19, 548–55 (1966).

21. Cui, Z. et al. Combining Sense and Nonsense Codon Reassignment for Site-Selective Protein Modification with Unnatural Amino Acids. ACS Synth. Biol. 6, 535–544 (2017).

22. d’Aquino, A. E., Kim, D. S. & Jewett, M. C. Engineered Ribosomes for Basic Science and Synthetic Biology. Annu. Rev. Chem. Biomol. Eng. 1–30 (2018). doi:10.1146/annurev-chembioeng-060817-084129

23. Danger, G., Plasson, R. & Pascal, R. Pathways for the formation and evolution of peptides in prebiotic environments. Chem. Soc. Rev. 41, 5416 (2012).

24. De Bodt, S., Maere, S. & Van de Peer, Y. Genome duplication and the origin of angiosperms. Trends Ecol. Evol. 20, 591–597 (2005).

25. Desai, M. M. & Fisher, D. S. Beneficial Mutation Selection Balance and the Effect of Linkage on Positive Selection. Genetics 176, 1759–1798 (2007).

26. Desai, M. M., Fisher, D. S. & Murray, A. W. The Speed of Evolution and Maintenance of Variation in Asexual Populations. Curr. Biol. 17, 385–394 (2007).

27. Endy, D. Foundations for engineering biology. Nature 438, 449–453 (2005).

28. Escudero, J. A. et al. Recoding of synonymous genes to expand evolutionary landscapes requires control of secondary structure affecting translation. Biotechnol. Bioeng. (2017). doi:10.1002/bit.26450

29. Fredens, J. et al. Total synthesis of Escherichia coli with a recoded genome. Nature 569, 514–518 (2019).

30. Fujishima, K. et al. Reconstruction of cysteine biosynthesis using engineered cysteine-free enzymes. Sci. Rep. 8, 1776 (2018).

31. Gallagher, R. R., Patel, J. R., Interiano, A. L., Rovner, A. J. & Isaacs, F. J. Multilayered genetic safeguards limit growth of microorganisms to defined environments. Nucleic Acids Res. 43, 1945–1954 (2015).

32. Gesteland, R., Weiss, R. & Atkins, J. Recoding: reprogrammed genetic decoding. Science (80-.). 257, 1640–1641 (1992).

33. Gorini, L., Jacoby, G. A. & Breckenridge, L. Ribosomal Ambiguity. Cold Spring Harb. Symp. Quant. Biol. 31, 657–64 (1966).

34. Gregory, S. T., Lieberman, K. R. & Dahlberg, A. E. Mutations in the peptidyl transferase region of E.coli 23s rRNA affecting translational accuracy. Nucleic Acids Res. 22, 279–284 (1994).

35. Harashima, F. Recent advances of mechatronics. in Proceedings of IEEE International Symposium on Industrial Electronics 1, 1–4 (IEEE, 1996).

36. Hershberg, R. & Petrov, D. A. Selection on Codon Bias. Annu. Rev. Genet. 42, 287–299 (2008).

37. Hinegardner, R. T. & Engelberg, J. Rationale for a Universal Genetic Code. Science (80-.). 142, 1083–1085 (1963).

38. House, T. W. National Bioeconomy Blueprint, April 2012. Ind. Biotechnol. 8, 97–102 (2012).

39. Johnson, D. B. F. et al. Release Factor One Is Nonessential in Escherichia coli. ACS Chem. Biol. 7, 1337–1344 (2012).

40. Kasahara, M. et al. Chromosomal localization of the proteasome Z subunit gene reveals an ancient chromosomal duplication involving the major histocompatibility complex. Proc. Natl. Acad. Sci. U. S. A. 93, 9096–101 (1996).

41. Katz, L. et al. Synthetic biology advances and applications in the biotechnology industry: a perspective. J. Ind. Microbiol. Biotechnol. 45, 449–461 (2018).

42. Keasling, J. D. Synthetic biology for synthetic chemistry. ACS Chem. Biol. 3, 64–76 (2008).

43. Khalil, A. S. & Collins, J. J. Synthetic biology: Applications come of age. Nat. Rev. Genet. 11, 367–379 (2010).

44. Kirsebom, L. A. & Isaksson, L. A. Involvement of ribosomal protein L7/L12 in control of translational accuracy. Proc. Natl. Acad. Sci. U. S. A. 82, 717–21 (1985).

45. Koonin, E. V. & Novozhilov, A. S. Origin and Evolution of the Universal Genetic Code. Annu. Rev. Genet. 51, 45–62 (2017).

46. Koonin, E. V. & Novozhilov, A. S. Origin and evolution of the genetic code: The universal enigma. IUBMB Life 61, 99–111 (2009).

47. Kugelberg, E., Kofoid, E., Reams, A. B., Andersson, D. I. & Roth, J. R. Multiple pathways of selected gene amplification during adaptive mutation. Proc. Natl. Acad. Sci. U. S. A. 103, 17319–24 (2006).

48. Kyte, J. & Doolittle, R. F. A simple method for displaying the hydropathic character of a protein. J. Mol. Biol. 157, 105–132 (1982).

49. Lajoie, M. J. et al. Genomically Recoded Organisms Expand Biological Functions. Science (80-.). 342, 357–360 (2013).

50. Lee, J. W., Chan, C. T. Y., Slomovic, S. & Collins, J. J. Next-generation biocontainment systems for engineered organisms. Nat. Chem. Biol. 1 (2018). doi:10.1038/s41589-018-0056-x

51. Loewe, L. & Hill, W. G. The population genetics of mutations: good, bad and indifferent. Philos. Trans. R. Soc. B Biol. Sci. 365, 1153–1167 (2010).

52. Lorenzini, D. M., da Silva, P. I., Fogaça, A. C., Bulet, P. & Daffre, S. Acanthoscurrin: a novel glycine-rich antimicrobial peptide constitutively expressed in the hemocytes of the spider Acanthoscurria gomesiana. Dev. Comp. Immunol. 27, 781–791 (2003).

53. Magliery, T. J., Anderson, J. C. & Schultz, P. G. Expanding the genetic code: selection of efficient suppressors of four-base codons and identification of “shifty” four-base codons with a library approach in Escherichia coli11Edited by M. Gottesman. J. Mol. Biol. 307, 755–769 (2001).

54. McClory, S. P., Devaraj, A. & Fredrick, K. Distinct functional classes of ram mutations in 16S rRNA. RNA 20, 496–504 (2014).

55. McClory, S. P., Leisring, J. M., Qin, D. & Fredrick, K. Missense suppressor mutations in 16S rRNA reveal the importance of helices h8 and h14 in aminoacyl-tRNA selection. RNA 16, 1925–34 (2010).

56. Miller, S. L. A Production of Amino Acids Under Possible Primitive Earth Conditions. Science (80-.). 117, 528–529 (1953).

57. Mittal, S. & Vetter, J. S. A Survey of Software Techniques for Using Non-Volatile Memories for Storage and Main Memory Systems. IEEE Trans. Parallel Distrib. Syst. 27, 1537–1550 (2016).

58. Molina, L., Ramos, C., Ronchel, M.-C., Molin, S. & Ramos, J. L. Construction of an efficient biologically contained pseudomonas putida strain and its survival in outdoor assays. Appl. Environ. Microbiol. 64, 2072–8 (1998).

59. Moratorio, G. et al. Attenuation of RNA viruses by redirecting their evolution in sequence space. Nat. Microbiol. 2, 17088 (2017).

60. Murgola, E. J. et al. Variety of nonsense suppressor phenotypes associated with mutational changes at conserved sites in Escherichia coli ribosomal RNA. Biochem. Cell Biol. 73, 925–31 (1995).

61. Neumann, H., Slusarczyk, A. L. & Chin, J. W. De Novo Generation of Mutually Orthogonal Aminoacyl-tRNA Synthetase/tRNA Pairs. J. Am. Chem. Soc. 132, 2142–2144 (2010).

62. Neumann, H., Wang, K., Davis, L., Garcia-Alai, M. & Chin, J. W. Encoding multiple unnatural amino acids via evolution of a quadruplet-decoding ribosome. Nature 464, 441–444 (2010).

63. NIH OSP. NIH Guidelines for Research Involving Recombinant or Synthetic Nucleic Acid Molecules (April 2016). (2016).

64. Niu, W., Schultz, P. G. & Guo, J. An Expanded Genetic Code in Mammalian Cells with a Functional Quadruplet Codon. ACS Chem. Biol. 8, 1640–1645 (2013).

65. Nye, C. Biohacker: Meet the people ‘hacking’ their bodies - BBC News. Bbc (2018). Available at: https://www.bbc.com/news/technology-46442519. (Accessed: 23rd January 2019)

66. O’Connor, M. & Dahlberg, A. E. The Involvement of Two Distinct Regions of 23 S Ribosomal RNA in tRNA Selection. J. Mol. Biol. 254, 838–847 (1995).

67. O’Connor, M., Thomas, C. L., Zimmermann, R. A. & Dahlberg, A. E. Decoding fidelity at the ribosomal A and P sites: Influence of mutations in three different regions of the decoding domain in 16S rRNA. Nucleic Acids Res. 25, 1185–1193 (1997).

68. Osawa, S. & Jukes, T. H. On Codon reassignment. J. Mol. Evol. 41, 247–249 (1995).

69. Ostrov, N. et al. Design, synthesis, and testing toward a 57-codon genome. Science (80-.). 353, 819–822 (2016).

70. Piepersberg, W., Böck, A. & Wittmann, H. G. Effect of different mutations in ribosomal protein S5 of Escherichia coli on translational fidelity. Mol. Gen. Genet. 140, 91–100 (1975).

71. Pines, G., Winkler, J. D., Pines, A. & Gill, R. T. Refactoring the Genetic Code for Increased Evolvability. MBio 8, e01654–17 (2017).

72. Ravikumar, A. & Liu, C. C. Biocontainment through reengineered genetic codes. ChemBioChem 16, 1149–1151 (2015).

73. Redford, K. H., Adams, W. & Mace, G. M. Synthetic Biology and Conservation of Nature: Wicked Problems and Wicked Solutions. PLoS Biol. 11, e1001530 (2013).

74. Ronchel, M. C. & Ramos, J. L. Dual System To Reinforce Biological Containment of Recombinant Bacteria Designed for Rhizoremediation. Appl. Environ. Microbiol. 67, 2649–2656 (2001).

75. Rosset, R. & Gorini, L. A ribosomal ambiguity mutation. J. Mol. Biol. 39, 95–112 (1969).

76. Rovner, A. J. et al. Recoded organisms engineered to depend on synthetic amino acids. Nature 518, 89–93 (2015).

77. Santer, U. V et al. A mutation at the universally conserved position 529 in Escherichia coli 16S rRNA creates a functional but highly error prone ribosome. RNA 1, 89–94 (1995).

78. Schultz, D. W. & Yarus, M. Transfer RNA Mutation and the Malleability of the Genetic Code. J. Mol. Biol. 235, 1377–1380 (1994).

79. Silver, D. et al. Mastering the game of Go with deep neural networks and tree search. Nature 529, 484–489 (2016).

80. Silver, D. et al. A general reinforcement learning algorithm that masters chess, shogi, and Go through self-play. Science 362, 1140–1144 (2018).

81. Sniegowski, P. D. & Gerrish, P. J. Beneficial mutations and the dynamics of adaptation in asexual populations. Philos. Trans. R. Soc. B Biol. Sci. 365, 1255–1263 (2010).

82. Steidler, L. et al. Biological containment of genetically modified Lactococcus lactis for intestinal delivery of human interleukin 10. Nat. Biotechnol. 21, 785–789 (2003).

83. Tuite, M. F. Transfer RNA in Decoding and the Wobble Hypothesis. in Encyclopedia of Life Sciences 1–7 (John Wiley & Sons, Ltd, 2001). doi:10.1038/npg.els.0001497

84. Von Neumann, J. & Burks, A. W. Theory of self-reproducing automata. (Urbana, University of Illinois Press, 1966).

85. Wang, K. et al. Defining synonymous codon compression schemes by genome recoding. Nature 539, 59–64 (2016).

86. Wang, K., Schmied, W. H. & Chin, J. W. Reprogramming the Genetic Code: From Triplet to Quadruplet Codes. Angew. Chemie Int. Ed. 51, 2288–2297 (2012).

87. Wang, L. & Schultz, P. G. Expanding the Genetic Code. Angew. Chemie Int. Ed. 44, 34–66 (2005).

88. Watanabe, K. & Osawa, S. tRNA Sequences and Variations in the Genetic Code. in tRNA: Structure, Biosynthesis, and Function (eds. Soll, D. & Rajbhandary, U. L.) 224–250 (ASM Press, 1995).

89. Wolfe, K. H. & Shields, D. C. Molecular evidence for an ancient duplication of the entire yeast genome. Nature 387, 708–713 (1997).

90. Wright, A. F. Genetic Variation: Polymorphisms and Mutations. in Encyclopedia of Life Sciences (John Wiley & Sons, Ltd, 2005). doi:10.1038/npg.els.0005005

91. Yang, F., Moss, L. G. & Phillips, G. N. The molecular structure of green fluorescent protein. Nat. Biotechnol. 14, 1246–1251 (1996).

92. Yokoyama, S. & Nishimura, S. Modified Nucleosides and Codon Recognition. in tRNA: Structure, Biosynthesis, and Function 207–223 (ASM Press, 1995).

93. Yona, A. H. et al. Chromosomal duplication is a transient evolutionary solution to stress. Proc. Natl. Acad. Sci. 109, 21010–21015 (2012).

