## Supplementary Materials for "Synthetic Genetic Codes Designed to Hinder Evolution"

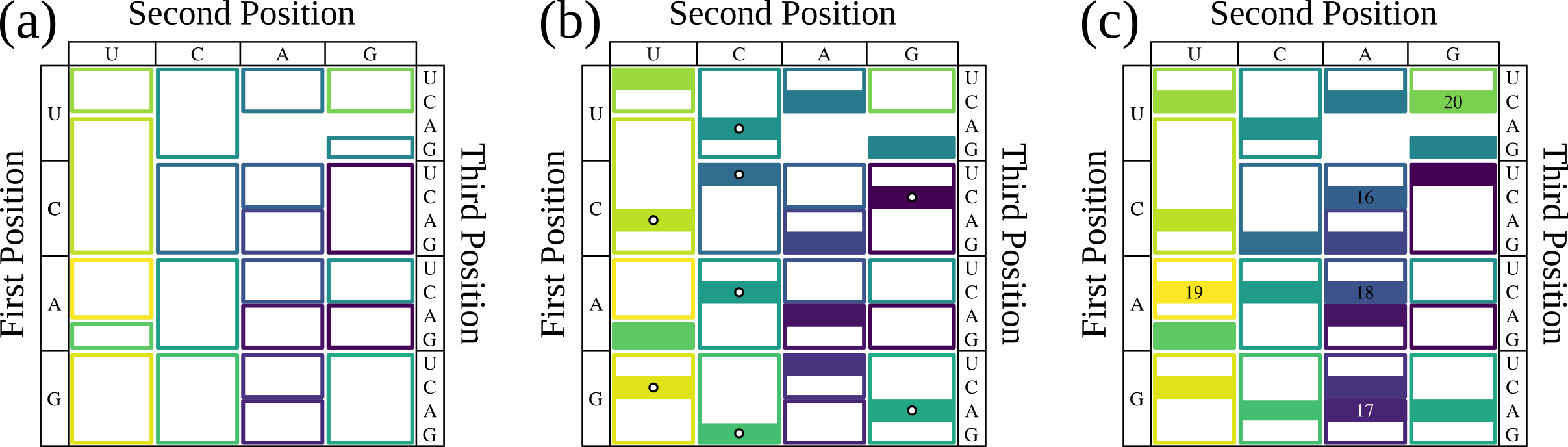
Supplementary Information

*Supplementary Figure 1: Constructing reduced amino acid set fail-safe genetic codes.* **(a)** Simplified representation of the Standard Code. Outlined boxes represent blocks of codons that encode the same amino acid. Color represents the rank ordered hydropathy of the encoded amino acid. **(b)** Construction of RED15. Due to the structure of the Standard Code, some pairs of amino acids cannot be simultaneously encoded such that their sense codons are not adjacent by point mutation. These cases occur when two amino acids co-occupy an NNX codon block. For example, aspartate is encoded by GAY and glutamate by GAR, where Y is a pyrimidine (U or C), and R is a purine (A or G). We first assigned amino acids not affected by this constraint (marked by white dots). There are seven amino acid pairs that conflict within an NNX block: phenylalanine (UUY) and leucine (UUR), isoleucine (AUH) and methionine (AUG), histidine (CAY) and glutamine (CAR), asparagine (AAY) and lysine (AAR), cysteine (UGY) and tryptophan (UGG), serine (AGY) and arginine (AGR), and aspartate and glutamate as mentioned previously. In two cases – F and L, and S and R – this conflict can be resolved by encoding one amino acid in another NNX block. As a result, only the following conflicting pairs of amino acids require resolution: I/M, H/Q, N/K, C/W, and D/E. We used the following biological and biochemical arguments to choose between some pairs of amino acids: M over I to avoid re-engineering translation initiation and due to the physicochemical similarity of isoleucine to leucine; K over N because lysine can form intramolecular salt links and act as a general base; Q over H because glutamine can both accept and donate hydrogen bonds, and to retain one amino acid with an amide side chain; and D over E because aspartate acts as a biosynthetic precursor for methionine, threonine, and lysine [Kanehisa and Goto 2000]. We chose W over C due to the following code structure argument: F (UUY) and Y (AUY) are constrained to two codons each such that either C codon (GUY) would be adjacent by point mutation to F or Y. W (UGG) is not similarly constrained, thus we chose W. We concede convincing arguments can be made for including different combinations of amino acids, but any resulting codes using codons that are nonadjacent by point mutation will have the same predicted evolutionary rate.

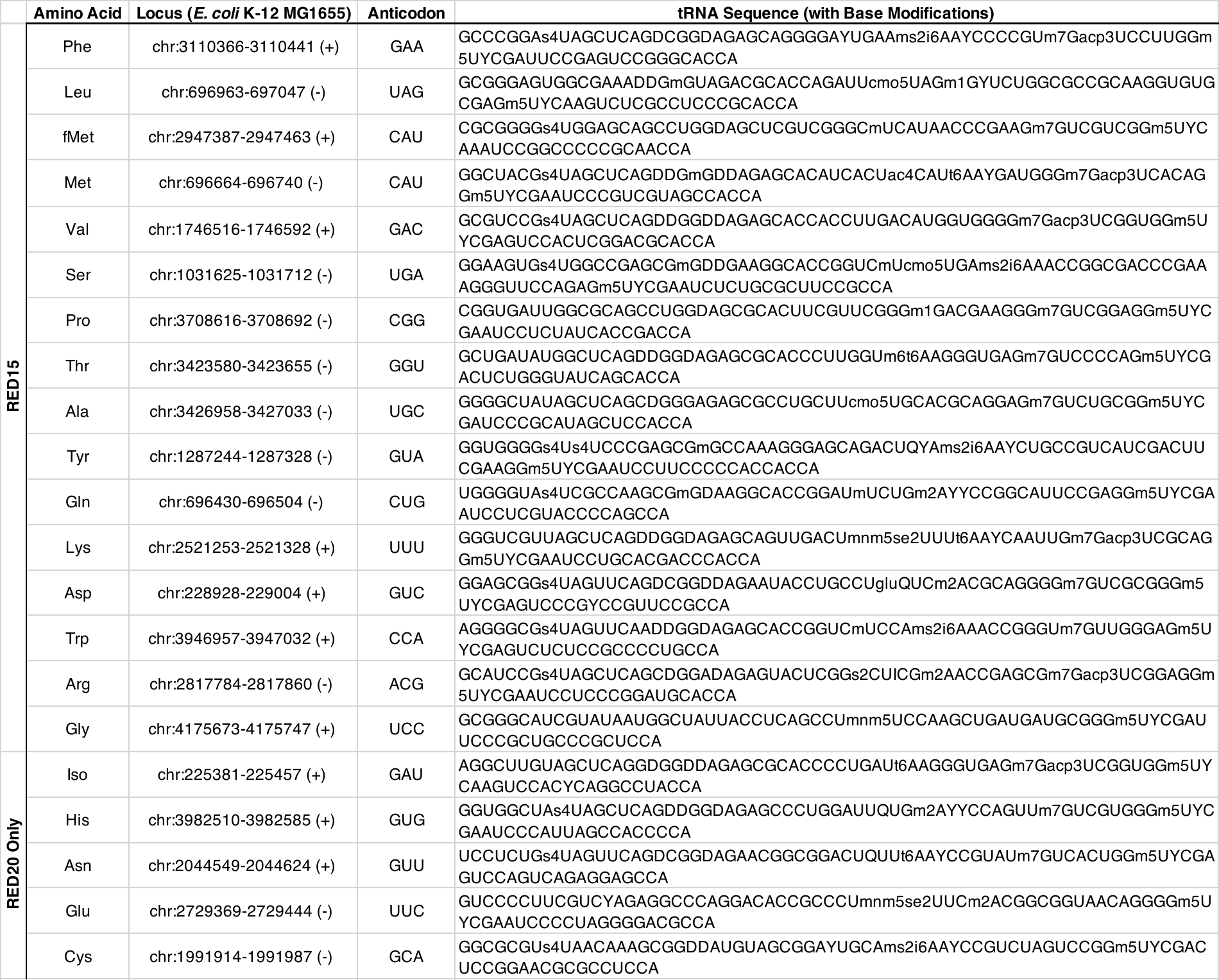

*Supplementary Table 1: Mature Sequences of tRNAs used in RED15 and RED20 genetic codes.* Proposed set of tRNAs for instantiating RED15 and RED20, along with their genomic locations in E. coli (K-12 MG1655) and their mature tRNA sequences with base modifications. Shorthand symbols for the base modifications used by these tRNA species: D, dihydrouridine; Y, pseudouridine; I, inosine; Q, queuosine; gluQ, glutamyl-queuosine; Um, 2’-o-methyluridine; m5U, 5-methyluridine; s4U, 4-thiouridine; mnm5U, 5-methylaminomethyluridine; mnm5s2U, 5-methylaminomethyl-2-thiouridine; cmo5U, uridine 5-oxyacetic acid; acp3U, 3-(3-amino-3-carboxypropyl)-uridine; Cm , 2’-o-methylcytidine; s2C, 2-thiocytidine; ac4C, N4-acetylcytidine; m2A, 2-methyladenosine; t6A, N6-threonylcarbamoyladenosine; m6t6A, N6-methyl-N6-threonylcarbamoyladenos­ine; ms2i6A, 2-methylthio-N6-isopentenyladenosine; Gm, 2’-o-methylguanosine; m1G, 1-methylguanosine; m7G, 7-methylguanosine. We require single base changes compared to wild type sequences for 5 tRNA species (proline, alanine, arginine, isoleucine, and cysteine), highlighted in red in each case. Genomic loci and sequences were taken from the GtRNAdb 2.0 database [Chan and Lowe 2016]. Base modifications were taken from the Modomics RNA modification database [Boccaletto et al. 2017].

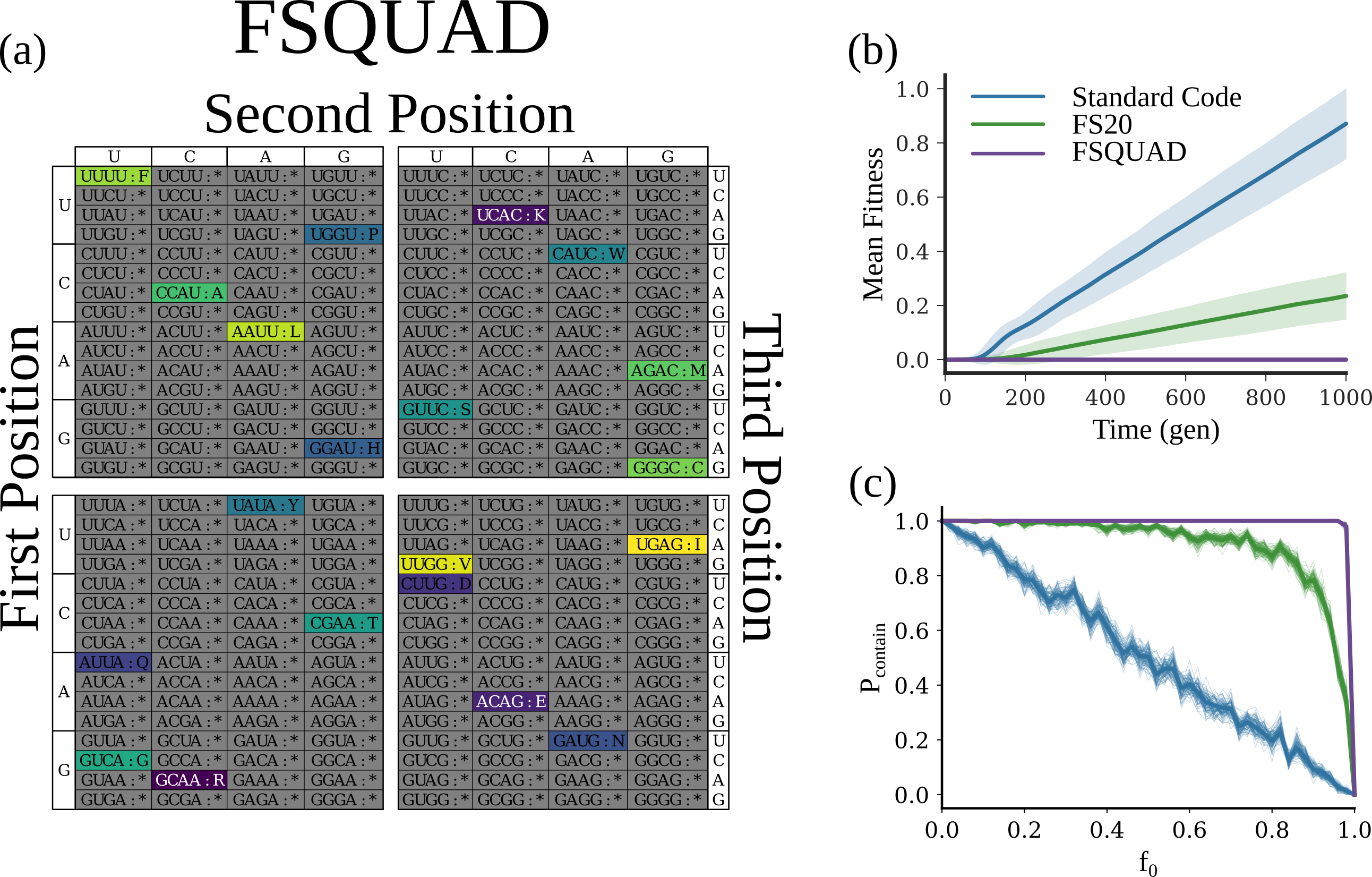
*Supplementary* *Figure 2: Quadruplet fail-safe genetic codes may support expression with 20 or more amino acids without permitting missense mutations.* **(a)** Table representation of a quadruplet fail-safe genetic code encoding all 20 proteinogenic amino acids (FSQUAD). Color signifies the rank-ordered hydropathy of the amino acids (I is most hydrophobic, R is most hydrophilic). Quadruplet decoding rather than triplet decoding allows for 256 (4^4^) sense codons rather than 64 (4^3^). As a result, FSQUAD has a set of 64 codons that all vary from each other at 2 or more positions, rather than the set of 16 available to FS20 or FS16. **(b)** Mean fitness traces for fail-safe codes using triplet and quadruplet-codon decoding (n = 1000 replicates). Bold lines indicate the mean fitness of a batch culture, averaged across replicates. Shaded regions represent the standard deviation of the mean fitness across replicates. The Standard Code is represented in blue. FS20 is represented in green. FSQUAD is represented in purple. **(c)** $P_{contain}$ at steady state vs. $f_{0}$ for invasive strains using fail-safe codes using triplet and quadruplet-codon decoding (n = 300 replicates). Colors are the same as in Fig 6b. Lighter shaded lines represent bootstrap-resampled traces of the data.

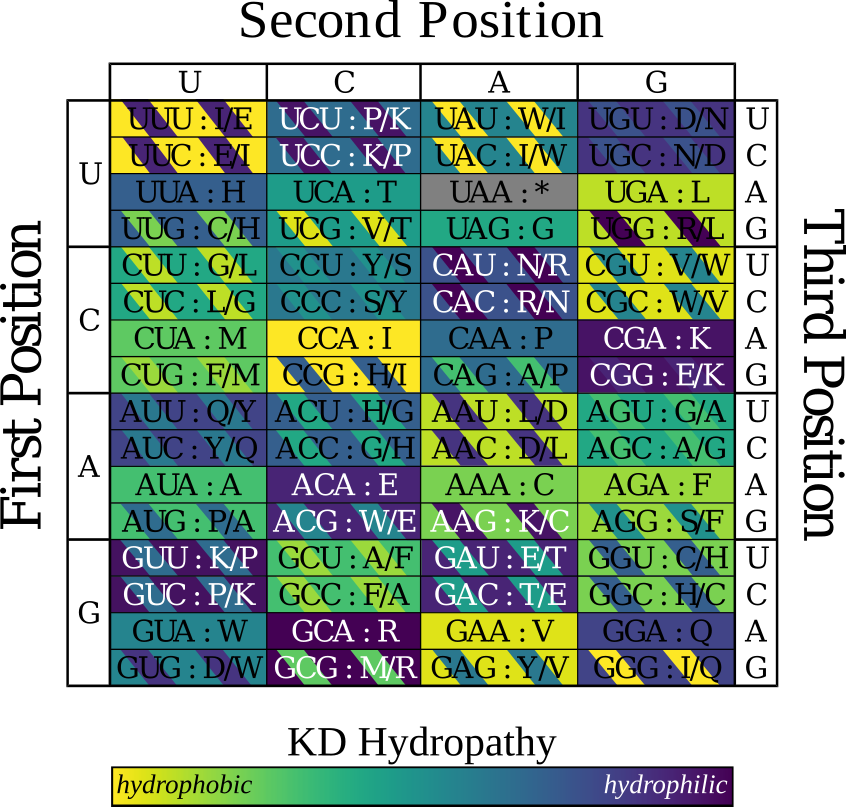

*Supplementary Figure 3: Predicted effect of tRNA promiscuity on Colorado Code.* The hyperevolvable code proposed by Pines et al. 2017, adjusted to consider ambiguous decoding due to tRNA promiscuity. Color signifies the rank-ordered hydropathy of the amino acids (I is most hydrophobic, R is most hydrophilic). Codons shaded with two colors would be recognized by two tRNA species, with each color corresponding to an amino acid decoded at that codon. For these ambiguous codons the first amino acid listed corresponds to the cognate tRNA, with the second amino acid corresponding to the wobble decoding tRNA (e.g., UUU is assigned to I, but also encodes E by wobble decoding). Only 17 of 64 codons are unambiguously decoded, one of which corresponding to the termination signal.

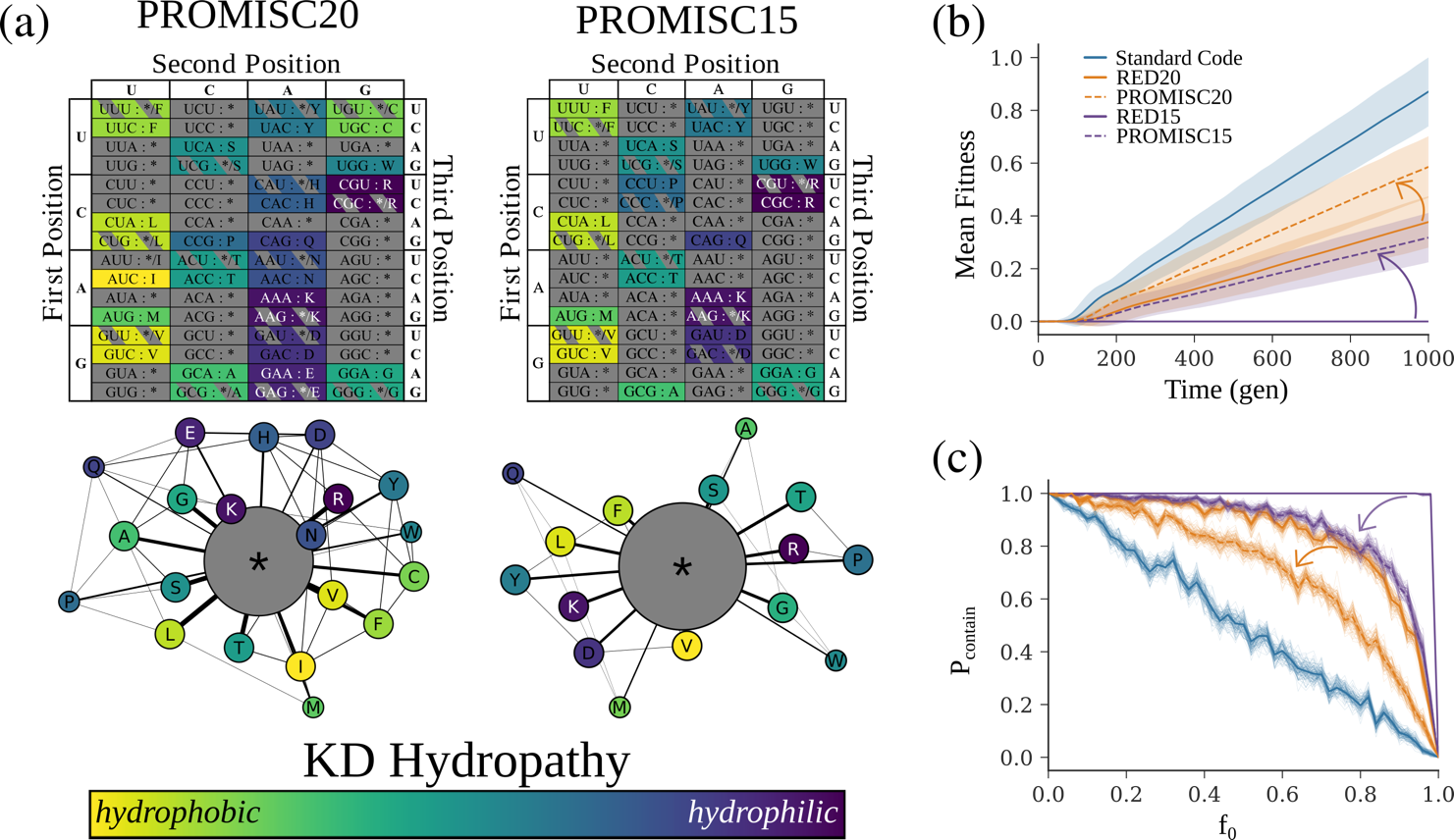

*Supplementary Figure 4: Accounting for tRNA promiscuity has a limited effect on the evolutionary rate of fail-safe codes.* **(a)** Effect of wobble decoding on RED20 and RED15. Table and mutation-distance network representations of the codes that result from considering wobble decoding (PROMISC20 resulting from RED20 and PROMISC15 resulting from RED15). Color signifies the rank-ordered hydropathy of the amino acids (I is most hydrophobic, R is most hydrophilic). Shaded boxes represent null codons that may be recognized by wobble decoding. A detailed discussion of tRNA promiscuous decoding is included in the Materials and Methods section. **(b)** Mean fitness traces for fail-safe codes with and without considering tRNA promiscuity (n = 1000 replicates). Bold lines indicate the mean fitness of a batch culture, averaged across replicates. Shaded regions represent the standard deviation of the mean fitness across replicates. The Standard Code is represented in blue. RED20 and RED15 are represented as the solid and dashed orange lines respectively. PROMISC20 and PROMISC15 are represented with solid and dashed purple lines respectively. **(c)** $P_{contain}$ at steady state vs. $f_{0}$ for invasive strains using fail-safe codes with and without considering wobble decoding (n = 300 replicates). Colors are the same as in Fig. 5b. Lighter shaded lines represent bootstrap-resampled traces of the data.

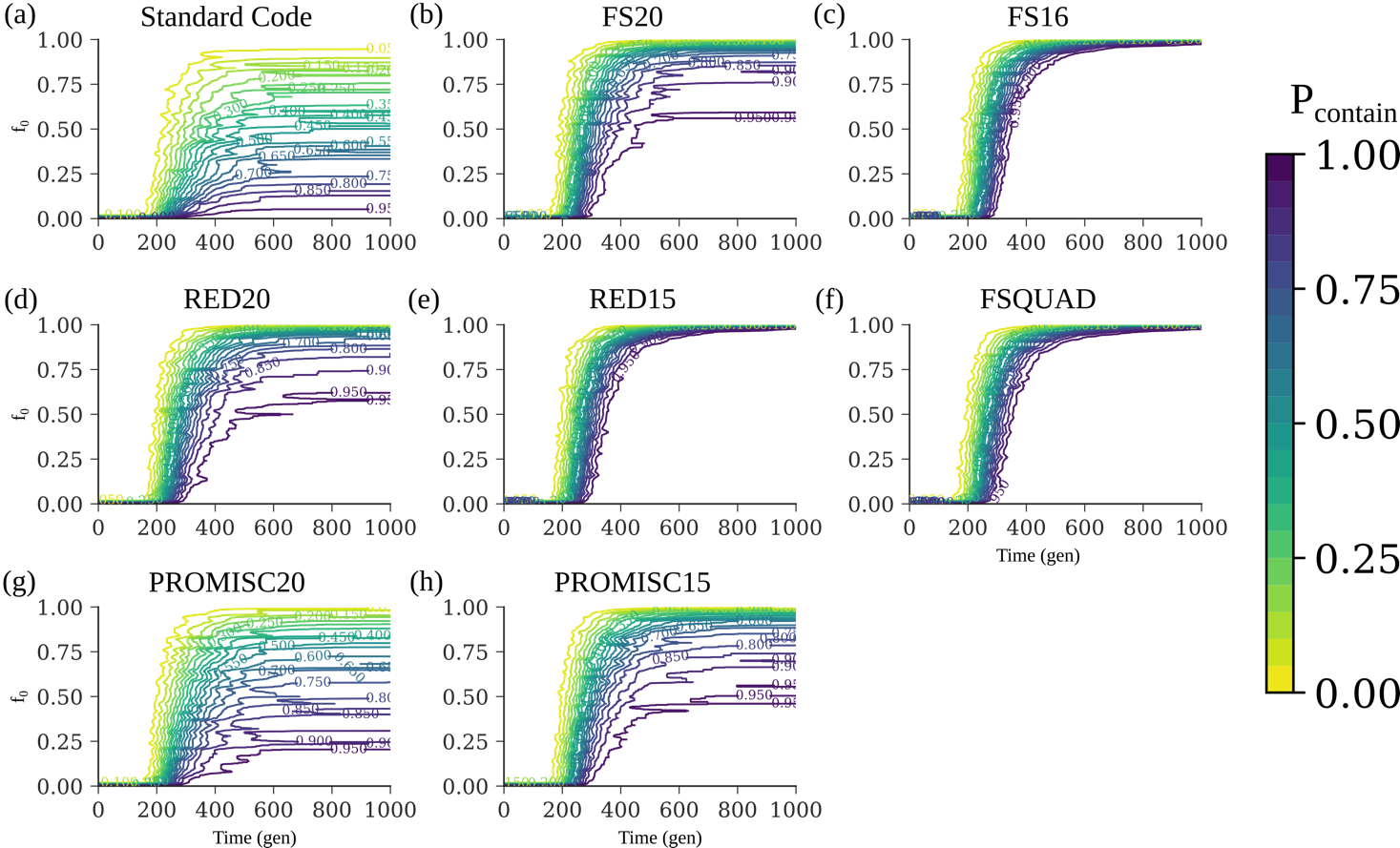

*Supplementary Figure 5: Containment probability vs. initial invasive population fraction and vs. time for fail-safe genetic codes.* Contour graphs of containment probability vs. time (x axis) and initial population fraction (y axis) for invasive strains using **(a)** the Standard Code, **(b)** FS20, **(c**) FS16, **(d)** RED20, **(e)** RED15, **(f)** FSQUAD, **(g)** PROMISC20, and **(h)** PROMISC15 (n = 300 replicates). Color represents magnitude, varying from 0 (yellow) to 1 (purple). $P_{contain}$reaches a steady state value at the limit of large *t*.

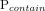

| **Position** | **Amino Acid** | **Conservation Score** | **MSA** | **Residue Diversity** |
| --- | --- | --- | --- | --- |
| 5 | E | -0.145 | 11/46 | {'P', 'A', 'E', 'R', 'K'} |
| 6 | E | -0.492 | 14/46 | {'L', 'A', 'Q', 'E', 'K'} |
| 14 | I | 0.602 | 23/46 | {'S', 'L', 'V', 'M', 'Y', 'C', 'F', 'T', 'I'} |
| 17 | E | 0.153 | 37/46 | {'N', 'S', 'H', 'F', 'D', 'E', 'R', 'T'} |
| 23 | N | -1.222 | 45/46 | {'N', 'G', 'H', 'D', 'E'} |
| 25 | H | -0.873 | 45/46 | {'N', 'H', 'M', 'Q', 'K'} |
| 32 | E | 1.616 | 45/46 | {'N', 'S', 'H', 'V', 'Q', 'D', 'E', 'K', 'T'} |
| 34 | E | 2.599 | 45/46 | {'N', 'S', 'L', 'V', 'A', 'Y', 'Q', 'D', 'E', 'K', 'T', 'I', 'R'} |
| 47 | I | 1.041 | 45/46 | {'N', 'S', 'V', 'C', 'Y', 'I', 'D', 'E', 'K', 'T', 'W', 'R'} |
| 48 | C | -1.12 | 45/46 | {'S', 'L', 'C', 'V', 'I'} |
| 70 | C | 0.599 | 46/46 | {'P', 'L', 'V', 'A', 'M', 'C', 'Q', 'F', 'I'} |
| 77 | H | 0.414 | 46/46 | {'N', 'S', 'G', 'H', 'V', 'D', 'E'} |
| **81** | **H** | **0.257** | 2/46 | **{'N', 'H'}** |
| 90 | E | -0.029 | 46/46 | {'S', 'V', 'A', 'M', 'D', 'E', 'K', 'T'} |
| 95 | E | -0.43 | 46/46 | {'H', 'Q', 'D', 'E', 'R', 'T'} |
| 98 | I | -0.138 | 46/46 | {'L', 'V', 'M', 'F', 'T', 'I'} |
| 105 | N | -0.348 | 46/46 | {'N', 'S', 'V', 'A', 'C', 'Q', 'F', 'K', 'T', 'I'} |
| 111 | E | 1.798 | 46/46 | {'N', 'H', 'V', 'A', 'M', 'Q', 'D', 'E', 'K', 'T', 'R'} |
| 115 | E | 1.472 | 46/46 | {'S', 'L', 'V', 'A', 'Q', 'D', 'E', 'K', 'T', 'I', 'R'} |
| 121 | N | -0.508 | 46/46 | {'N', 'S', 'H', 'V', 'A', 'Y', 'Q', 'D'} |
| 123 | I | -0.936 | 46/46 | {'S', 'V', 'F', 'T', 'I'} |
| 124 | E | 0.671 | 46/46 | {'N', 'S', 'M', 'I', 'Q', 'D', 'E', 'K', 'T', 'R'} |
| 128 | I | 0.061 | 46/46 | {'N', 'S', 'G', 'L', 'V', 'M', 'Q', 'D', 'E', 'K', 'T', 'I'} |
| 132 | E | 1.541 | 46/46 | {'S', 'P', 'H', 'L', 'V', 'A', 'I', 'D', 'E', 'K', 'T', 'R'} |
| 135 | N | -1.111 | 46/46 | {'N', 'V', 'P', 'H'} |
| 136 | I | -1.148 | 46/46 | {'L', 'V', 'T', 'I'} |
| 139 | H | -0.344 | 46/46 | {'N', 'S', 'H', 'M', 'Q', 'D', 'R', 'K'} |
| 142 | E | 2.252 | 46/46 | {'H', 'L', 'V', 'A', 'Y', 'M', 'D', 'E', 'T', 'I'} |
| 144 | N | 0.379 | 46/46 | {'N', 'S', 'H', 'G', 'M', 'Q', 'D', 'E', 'K', 'R'} |
| 146 | N | -0.378 | 46/46 | {'N', 'P', 'H', 'L', 'Q', 'D', 'E', 'R', 'T'} |
| 148 | H | 0.349 | 16/46 | {'N', 'S', 'H', 'G', 'V', 'Q', 'D'} |
| 149 | N | 0.611 | 46/46 | {'N', 'S', 'L', 'V', 'A', 'Y', 'M', 'Q', 'F', 'C', 'E', 'T', 'I'} |
| 152 | I | -0.405 | 46/46 | {'L', 'V', 'A', 'M', 'T', 'I'} |
| 159 | N | 0.157 | 46/46 | {'N', 'S', 'H', 'G', 'D', 'E', 'K', 'T'} |
| 161 | I | -0.707 | 46/46 | {'L', 'V', 'M', 'I'} |
| 164 | N | 0.72 | 46/46 | {'N', 'S', 'H', 'L', 'V', 'A', 'Y', 'F', 'D', 'E', 'I'} |
| 167 | I | -0.289 | 46/46 | {'N', 'V', 'A', 'Y', 'M', 'Q', 'F', 'I', 'K', 'R', 'W'} |
| 169 | H | -0.241 | 46/46 | {'H', 'Y', 'L', 'F'} |
| 170 | N | 0.677 | 46/46 | {'N', 'S', 'P', 'L', 'V', 'M', 'Q', 'D', 'E', 'K', 'T', 'R'} |
| 171 | I | -0.591 | 46/46 | {'L', 'V', 'K', 'T', 'I'} |
| 172 | E | 0.613 | 46/46 | {'N', 'V', 'A', 'M', 'Q', 'E', 'K', 'T'} |
| 181 | H | -0.432 | 46/46 | {'N', 'S', 'H', 'C', 'V', 'Q', 'F', 'T', 'I'} |
| 185 | N | 0.026 | 46/46 | {'N', 'H', 'G', 'L', 'V', 'A', 'M', 'E', 'K', 'R'} |
| 188 | I | 0.947 | 23/46 | {'N', 'G', 'L', 'V', 'D', 'I'} |
| 198 | N | 1.292 | 45/46 | {'N', 'S', 'P', 'V', 'A', 'Y', 'F', 'T', 'W'} |
| 199 | H | -1.559 | 45/46 | {'V', 'Y', 'H'} |
| 212 | N | 1.277 | 39/46 | {'N', 'S', 'L', 'Q', 'D', 'E', 'K', 'T', 'R'} |
| 213 | E | -0.907 | 37/46 | {'D', 'E', 'A', 'S'} |
| 217 | H | -0.337 | 27/46 | {'N', 'H', 'L', 'V', 'F', 'D', 'K'} |
| 222 | E | -0.597 | 25/46 | {'H', 'D', 'E', 'R', 'K'} |
| 229 | I | 0.228 | 7/46 | {'H', 'V', 'A', 'T', 'I'} |
| 231 | H | -0.519 | 4/46 | {'G', 'H'} |
| 235 | E | 0.318 | 3/46 | {'L', 'E'} |

*Supplementary Table 2: ConSurf analysis of residue conservation in fluorescent variants of GFP.* Positions in GFP encoding amino acids omitted from RED15 including position number (Position), consensus amino acid (Amino Acid), ConSurf conservation score (Conservation Score), number of variants in the multiple sequence alignment with an amino acid at the listed position (MSA), and all amino acids found at that position across fluorescent GFP variants (Residue Diversity) [Ashkenazy *et al.* 2016]. All positions show a diversity of encoded amino acids across fluorescent variants which include at least one amino acid in the RED20 supported amino acid set, except for except 81H (bolded). 81H however is only found in two of 46 variants in the multiple sequence alignment.

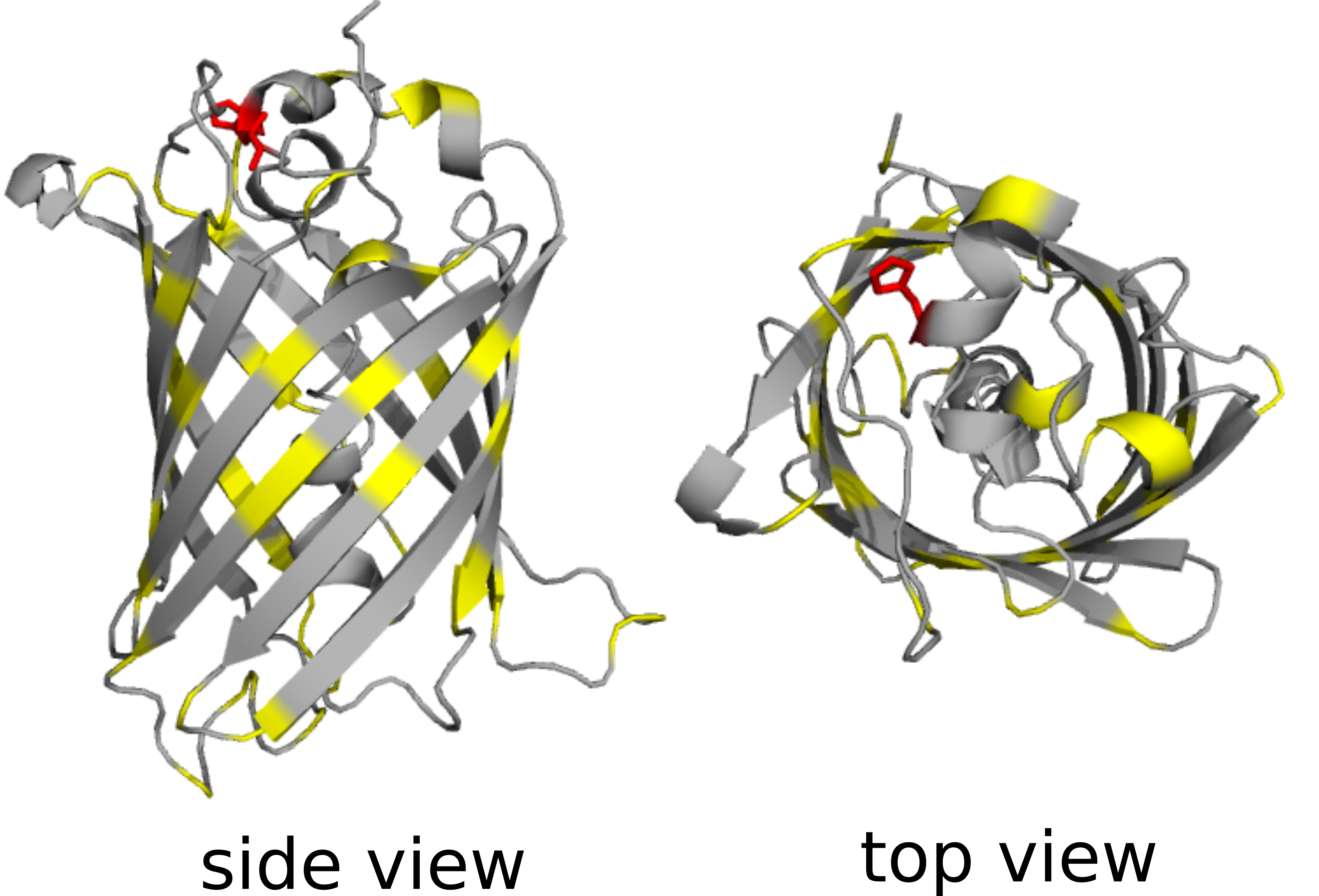

*Supplementary Figure 6: GFP structure with positions using non-RED20 amino acids highlighted.* Side (left) and top-down (right) views of GFP crystal structure (PDB: 1gfl) [Yang, Moss, and Phillips Jr. 1996]. Positions in which the canonically encoded amino acid is omitted from RED15 are marked in yellow. H81, the only position for which the residue diversity at the position lacks any amino acid in RED15, is marked in red.
